# A Quasi-Stationary Distribution Bound for Fault Analysis in Gene Regulatory Networks

**DOI:** 10.1101/2025.10.21.683707

**Authors:** Fabricio Cravo, Matthias Függer, Thomas Nowak

## Abstract

The inherent stochastic fluctuations in signaling molecules of Gene Regulatory Networks (GRNs) add unpredictability, complicating the design of robust synthetic GRNs that must function within precise ranges. Multi-stable GRNs, such as toggle switches, are central to systems like biosensors and logic gates but can fail due to unintended transitions between stable states caused by the fluctuations. Despite their importance, tools to characterize the probability distributions around stable states remain limited. We present a mathematical framework to analyze these multi-stable systems using continuous-time Markov chains (CTMCs) and quasi-stationary distributions. This framework is broadly applicable, requiring only that the state space is connected, making it applicable to a variety of systems. We then apply the framework to current examples from the literature and conclude that our method provides quantitative design principles for toggle switch design that match current experimental insights, identifying parameter thresholds where systems transition from frequent stochastic switching (hours) to stable operation (years to decades) and demonstrate upper bound calculations for false positive/negative rates in population-level biosensor dynamics.

## 1 Introduction

Gene Regulatory Networks (GRNs) are networks of genes that interact through signaling molecules [12]. These signaling molecules are often subject to stochastic fluctuations in their counts [36], which add unpredictability to their behavior. This unpredictability poses challenges in Synthetic Biology, the field focused on the rational engineering of life [6, 7]. Synthetic biologists design GRNs to engineer new cellular functionalities, but ensuring these systems function reliably within specific parameters is critical [27].

One class of such systems is multi-stable GRNs, where molecular signals govern gene expression with multiple stable states [12]. A well-known example of a multi-stable system is the toggle switch [19], a bi-stable GRN. Synthetic biologists design these switches to create cells capable of functioning as biological sensors, logic gates, and diagnostic tools [37, 42, 23, 5]. These systems are typically engineered to exhibit programmed transitions between stable states, triggered by environmental changes. However, the inherent stochastic fluctuations in biological systems give rise to non-negligible probabilities of transitions occurring between stable states even in the absence of the planned environmental triggers (i.e., stochastic switching).

Although preventing these unwanted transitions is critical for the proper functioning of synthetically engineered GRNs, methods to compute and bound the probability distributions around these stable states are lacking. Several publications informally describe these distributions as quasi-stationary distributions (QSDs) [38, 13]. QSDs are similar to stationary distributions, which are probability distributions that remain time-invariant, and in some biological systems, they are the long-term limit distribution [40]. Before converging to the stationary distribution, multi-stable GRNs exhibit distributions about each stable state that remain nearly unchanged for long periods before eventually converging [38, 13]. This long-lived, nearly invariant distribution is informally referred to as a QSD.

QSDs are well-defined mathematically in systems with absorbing states (i.e., states the system cannot leave once reached)[16, 34, 43], and several theorems simplify their calculation and analysis in such systems [43]. Formally, a QSD is a distribution that remains time-invariant provided the system has never reached an absorbing state. There are theorems guaranteeing the uniqueness of the QSD and that the system will converge under the condition that it cannot reach the absorbing states, if all states, except the absorbing ones, are connected to each other. Additionally, systems with absorbing states have a decay parameter that gives the probability of absorption by a given time.

Multi-stable GRNs can exhibit QSD-like behavior without absorbing states [3, 38, 26, 4]. Consequently, these systems lack the tools and theorems available for systems with absorption. Some researchers estimate transition times between stable states using the expected passage time, neglecting its variance [26, 3, 22, 38]. However, designing GRNs for applications where the timescale is shorter than this passage time is insufficient to guarantee proper functioning. If the variance is high enough, there may be a non-negligible probability that the network will transition between states unexpectedly [39]. This issue can be avoided with a direct estimation of transition probabilities.

Furthermore, some of the expected passage time calculations are designed only for bi-stable GRNs [3, 26], or they have restrictive assumptions about the model [3, 38]. For instance, Assaf et al. [3] assumed a large separation between stable states, a statement not necessarily true particularly when stochastic models are used for GRNs with low molecular counts [28]. Roma et al. [38] assumed that the distribution is quasi-stationary in the region of interest, a limitation when the QSD is not well understood.

In this paper, we provide a mathematical framework for analyzing QSDs in multi-stable GRNs without absorbing states. Our framework is built upon the Biochemical Reaction Network (BCRN) theory [18], which is akin to well-established Chemical Reaction Networks (CRNs) [15]. BCRNs are sets of species and reactions among the species [2], where the states correspond to counts of each of the species (for example, the state where counts are *A* = 10, *B* = 20, *C* = 5), and reactions represent state transitions (for example, the reaction *A* + *B* → *C* where the counts *A* and *B* decrease by one and *C* increases by one). Furthermore, BCRNs use Continuous Time Markov Chain (CTMC) to model the stochastic transitions between states, using reaction rates that define the transition probabilities. In this BCRN framework, we can analyze QSDs as probability distributions over species count configurations that remain approximately invariant over long time periods.

Our BCRN-based framework is generic, requiring only that the state space of the CTMC is connected, meaning all states can be reached from one another. From the original BCRN modeling the GRN, we construct what we call “cut-off process”. We define an equivalent CTMC on a subset of states *V* where all transitions that would lead outside *V* are instead redirected to a new absorbing state *a*. Within *V*, the cut-off process retains the original transition rates and dynamics.

This construction enables us to estimate the QSD by taking the limit of the conditional probability that the process remains within *V* as time approaches infinity. Additionally, we can calculate a decay parameter *λ* and a bound parameter *A* (which compensates for initial distribution effects), which together provide upper bounds on the probability of leaving *V*. When *V* encapsulates the region around a stable state, these parameters directly estimate the likelihood of unwanted transitions to other stable states.

We calculate probability bounds using this framework to several BCRN models from the literature to demonstrate its broad applicability and practical value. First, we analyze an Allee-based algorithm [9] for rare event detection, using our framework to estimate false positive and false negative rates by calculating the probability of unwanted transitions between detection states, enabling systematic parameter optimization for biosensor applications. Second, we conduct an analysis of bi-stable toggle switches across different promoter copy numbers (*P* ∈ *{*2, 6, 10, 20, 40*}*) and Hill coefficients (*n*_*h*_ ∈ {2, 3, 4}), identifying quantitative design principles that minimize stochastic switching and validating our theoretical predictions against established experimental design guidelines that successful toggle switches typically require more than 10 promoter copies. Finally, we apply our method to Barbier et al.’s [4] spatiotemporal pattern formation GRN, demonstrating the framework’s ability to analyze more models from literature.

Furthermore, we highlight the importance of accounting for the initial state probability distribution’s impact on the QSD. We demonstrate that different initial distributions in the vicinity of the same stable state for the same GRN can lead to substantially different QSDs. For practical application, we also provide a GitHub repository (github.com/BioDisCo/QSB_Bound_Algo) that allows users to input a BCRN in the MobsPy language [10] and a user-defined set *V* to calculate estimates of the QSD, decay parameter, and *A*.

## 2 Results

### 2.1 Model and Algorithm

The sequence of states of a process **X** = (*X*_0_, *X*_1_, *X*_2_, *X*_3_, …) is a sequence of tuples with each element of the sequence following 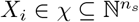, with *n*_*s*_ ∈ ℕ being the number of species in the BCRN. For each state *X*_*i*_, the elements of the tuple *X*_*i*_ are the available counts of a respective BCRN species (For an example see Figure 1a). The notation **X**(*t*) with *t* ∈ ℝ^+^, indicates the state of **X** at time *t*. Let the state space *χ* be connected for process **X**, which means that for every two states *x, y* ∈ *χ*, state *y* can be reached from state *x*. The matrix *Q* is the generator matrix of the process **X**, with *Q*(*x, y*) being the transition rate from state *x* to state *y* if *x* ≠ *y* and − Σ_*z*∈*X*_ *Q*(*x, z*) if *x* = *y*. Let *d* be an initial distribution for process **X**. If *d* is an initial distribution for process **X** and *A* an event, we use the notation

**Figure 1.**
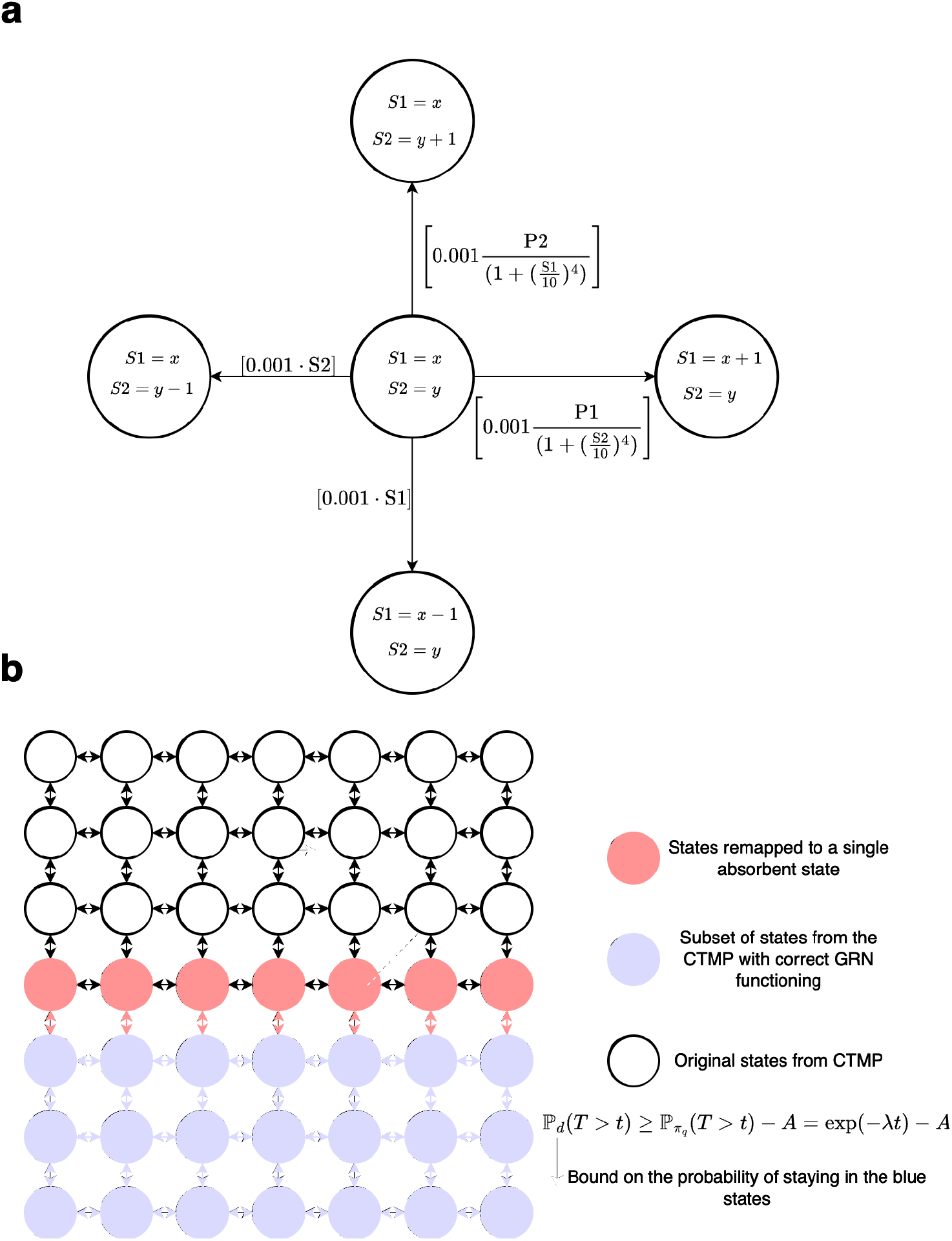
Illustration of the state-space for a cut-off process: **a** Example of GRN states in a Markov space using the toggle switch GRN from Equation (17). **b** Example of subset and equivalent absorbing state transitions for a cut-off process in *V*. The cut-off process cannot reach the states in white. The states in red are all remapped to transition to an absorbing state.

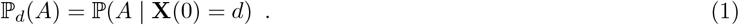

To predict if the state probability distribution of **X** approaches the QSD, we propose another continuous-time Markov process 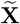 whose states are limited to a finite connected subset *V* ⊂ *χ* and an absorbing state *a* ∉ *V*. We refer to 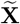 as the cut-off process in *V* as the reachable states of 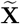 are a subset (“cut”) of the state space of **X**. By artificially creating an absorbing state *a*, we can apply the well-developed theory of QSDs with absorption to systems that originally had no absorbing states. This cut-out process construction allows us to estimate how long the original system stays near a stable state if the subset *V* compasses a region around it.

We also refer to **X** as the original process (original BCRN). Similarly to process **X**, denote by 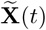 the state of 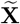 at time *t* and 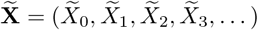 its sequence of states.

The transition rates from the generator matrix 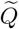 of 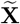 are equal to those of **X** for transitions between states within *V*. Any transition rate leading outside *V* leads to the absorbing state *a*. Let *x, y* ∈ *V*, *z* ∈ *χ*, and *a* ∈ *X\V*. The definition of 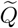 follows:

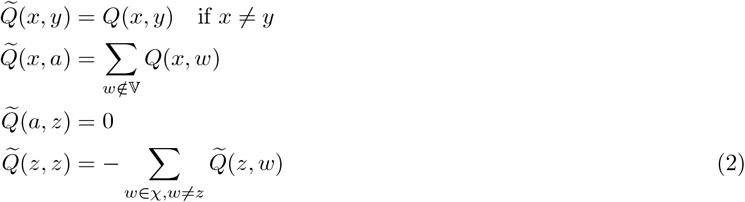

An example of the state spaces of a process **X** and a cut-out process 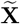 in *V* can be found in Figure 1b.

To define the QSD, we will use two random variables *T* and 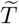, the time processes **X** and 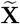 exit the set *V* respectively:

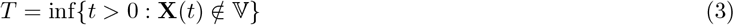

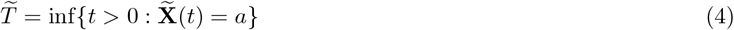

We refer to *T* and 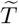 as exit times.

Denote *π*_*q*_ as the QSD of process 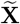. The QSD of process 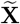 fulfills:

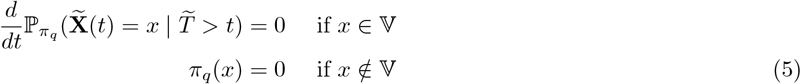

Furthermore, let *d*_*V*_ be an initial distribution concentrated on *V*, meaning *d*_V_(*x*) > 0 only for states *x* ∈ *V*. We prove in Appendix B that the following limit yields the same QSD from Equation (5):

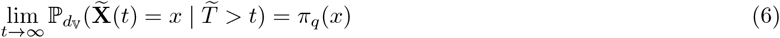

In Appendix B.1, we prove that *T* and 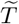 are equal in distribution. We also prove the following:

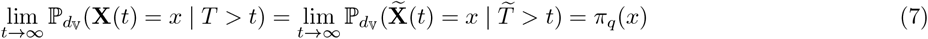

It follows from the law of total probability applied to process **X**:

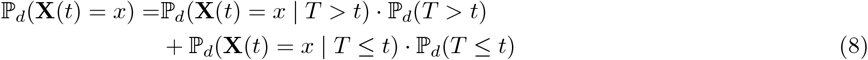

If we had ℙ_*d*_(*T* ≤ *t*) = 0 for all *t ≥* 0, then from Equation (7), ℙ_*d*_(**X**(*t*) = *x*) converges to *π*_*q*_(*x*). However, for connected Markov processes, the assumption that ℙ_*d*_(*T ≤ t*) = 0 does not hold in general. In Section 2.3, we take a numerical approach. We will show, using examples, that for some choices of subset *V* the probability ℙ_*d*_(*T ≤ t*) negligible and the process **X** approaches the QSD of the original BCRN before converging to the stationary distribution of **X** in a much larger time scale.

According to Van Doorn et al. [43], if the initial distribution for the process 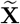 is the QSD *π*_*q*_, then:

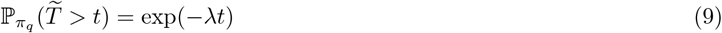

where 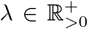 is called the decay parameter of generator matrix 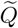. Further, from the equality in distribution of *T* and 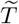, Equation (9) holds for *T* as well. The decay parameter gives the probability of non-absorption until time *t* given the QSD *π*_*q*_ of the cut-off process as an initial distribution. The parameter is always positive and is the negative of the non-zero eigenvalue of 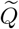 with the greatest real part [43].

To estimate if process **X** will approach the quasi-stationary distribution, we develop two complementary probability bounds: a general bound (Theorem 1) applicable to any connected CTMC using Algorithm 1 for a compensatory constant *A*, and a specialized bound for bounded birth-death processes (Section 2.2) that eliminates the need for a compensatory term for specific initial distributions.

The general bound uses the jump chain *J* from the generator matrix 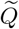, where *J* (*x, y*) gives the probability that the next state is *y* given the current state is *x*:

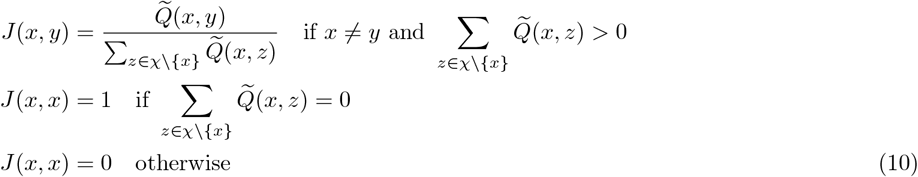

In Algorithm 1, the function `calculate_jump_chain` 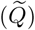 computes this jump chain from the generator matrix.

The correctness of the general bound with a compensatory constant *A* is stated in Theorem 1. For the proof, we refer the reader to Appendix B.

#### Theorem 1

(General QSD Bound). *Let* **X** *be a connected CTMC on totally ordered state space χ with generator matrix Q. For the cut-off process* 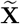 *with generator matrix* 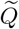 *on V* ∪*{a} where V* ⊂ *X is finite*, *V is connected for* 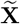, *and a* ∈*/ V*, *Algorithm 1 produces a constant A* ∈ ℝ_>0_ *such that for any initial distribution d:*

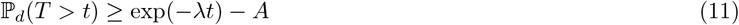

*where λ* ∈ ℝ_>0_ *is the decay parameter of* 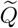 *and T* = inf*{t* > 0 : **X**(*t*) ∉ *V}*.

Algorithm 1 produces a constant *A* that serves as a compensatory term when the initial distribution *d* differs from the QSD *π*_*q*_. The constant *A* increases particularly when the initial distribution concentrates probability near the boundary states of *V* since these states have a higher exit probability.

### 2.2 QSD Bound for Bounded Birth-Death Markov Processes

The second bound that does not require the compensatory constant *A* applies to birth-death Markov processes. A birth-death Markov process is one whose state transitions can only be births that increase an integer state value by one or deaths that decrease it by one. Furthermore, if a birth-death Markov process is bounded, its state value has a maximum. If **X** is a bounded birth-death Markov process, there are initial distributions for which the QSD can bound the exit times without requiring the constant *A* from Algorithm 1, Let *d*_*b*_ be one of these distributions. It follows:

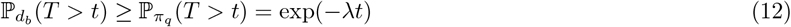

#### Algorithm 1

Quasi-Stationary Bound Algorithm

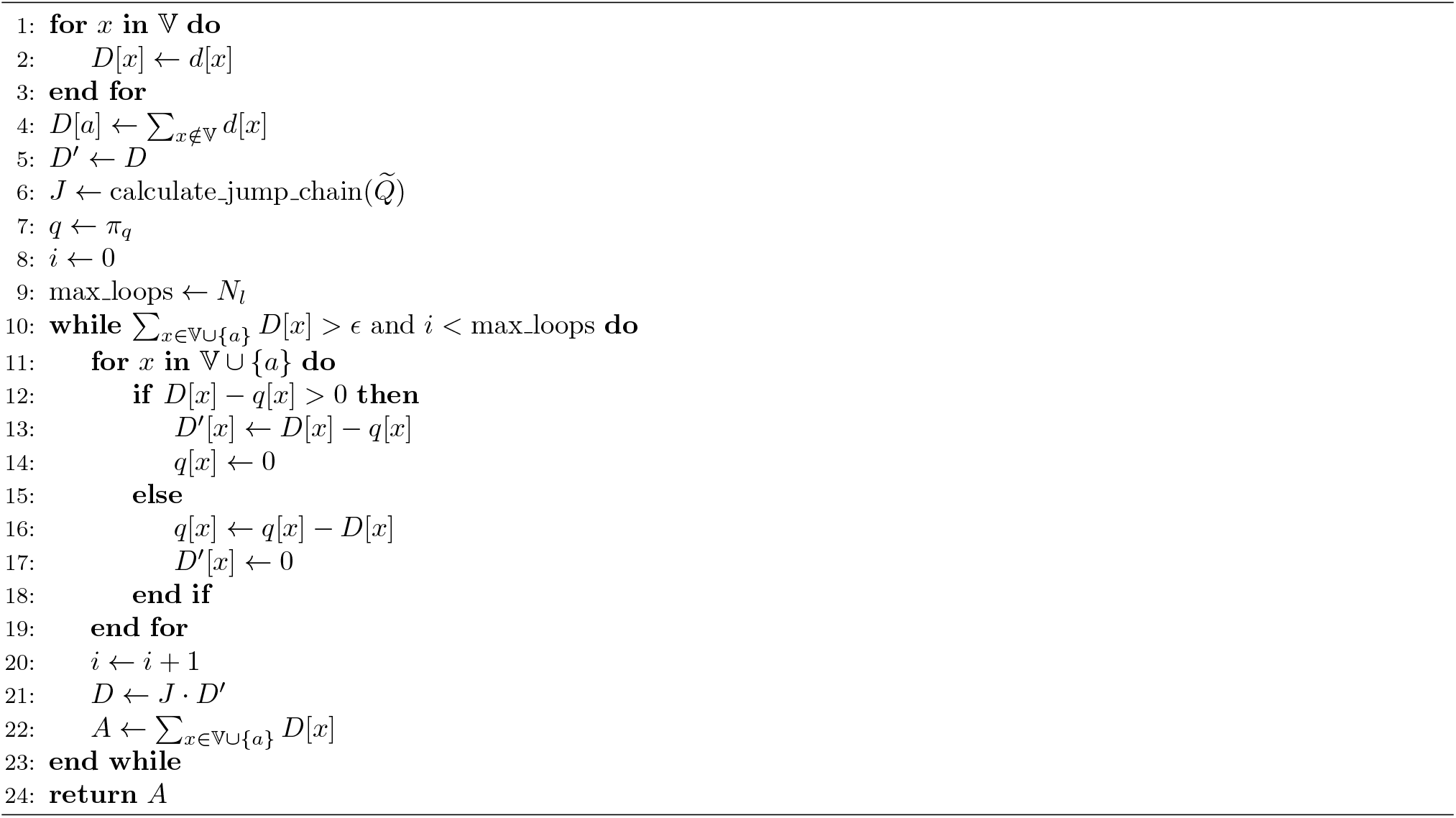

The exit time probability can be bound directly from estimating the decay parameter from the eigenvalues of the generator matrix 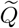 from the cut-off process, or by solving Equation (9) numerically. This results in a better and faster estimation of the bound when compared to the more generic approach from Theorem 1.

Theorem 2 establishes sufficient conditions for birth-death processes where the exit time probability is bounded by the quasi-stationary distribution without requiring the compensatory constant. It exploits the monotonicity property of birth-death processes: without loss of generality, exit probabilities decrease with distance from the absorbing boundary. The theorem uses auxiliary functions *v*_*y*_(*w*) that redistribute probability mass from the QSD *π*_*q*_ within the cut-off region *V* Each auxiliary function *v*_*y*_ aggregates probability from states *w ≤ y* (closer to the exit boundary than *y*). The first constraint requires that for each state *y*:

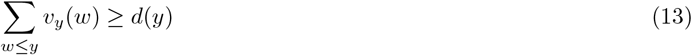

with *v*_*y*_(*w*) = 0 for *w* > *y*. The second constraint ensures the total redistributed mass does not exceed the QSD:

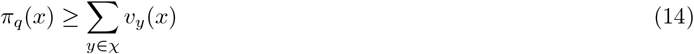

Together, these constraints guarantee that the initial distribution *d* has a lower exit probability than the QSD *π*_*q*_ for the cut-off process in *V*.

The second approach’s correctness is stated in Theorem 2, with the proof in Appendix B.3:

#### Theorem 2

(Birth-Death Process Bound). *Let* **X** *be a birth-death CTMC on χ* = ℕ *with cut-off process* 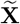 *on V* ∪ *{a} where V* ⊂ *X is finite and connected for* 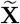, *and a* ∈*/ V*. *Without loss of generality, assume a* < *x for all x ∈ V*. *If an initial distribution d satisfies the following conditions:*

1. *For each state y* ∈ *χ, there exist functions v*_*y*_ : *χ*→ ℝ^+^ *such that* Σ_*w*≤*y*_ *v*_*y*_(*w*) ≥ *d*(*y*)
2. *For the QSD π*_*q*_ *of the cut-off process: π*_*q*_(*x*) ≥ *Σ*_*y*∈*X*_*v*_*y*_(*x*) *for all x* ∈ *X*

*then* ℙ_*d*_(*T* > *t*) ≥ exp(−*λt*), *where λ* ∈ ℝ_>0_ *is the decay parameter of* 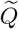 *and T* = inf{*t* > 0 : **X**(*t*) ∈*/ V*}.

We use Theorem 2 to perform the validation of a stochastic model of a rare event detection algorithm presented in our previous work [9] through Proposition 1. In this proposition, we propose two cut-off processes instead of one. One cut-out process created using the connected set *V*_*L*_ ⊂ *χ* of the lowest states in from the state space of the original process *χ* and an *a*_*L*_ greater than any state in *V*_*L*_. The other cut-off process is defined similarly, but it instead uses the greatest states of *χ* for its state space *V*_*H*_ ⊂ *χ*, and its absorbing state *a*_*H*_ is lower than every state in *V*_*H*_. Consequently, there are two quasi-stationary distributions, one for the cut-out process in *V*_*L*_ and the other for *V*_*H*_ that bound the exit times of their respective sets.

Proposition 1 provides a specific application of Theorem 2 for bounded birth-death chains. The proposition focuses on boundary initial distribution starting at the extreme states 0 or *P*, where *P* is the largest state achieved by the chain and analyzes the probability of transitions between opposite boundary regions. For the rare event detection context, this corresponds to studying false positive rates (unwanted transitions from an analyte-absent region) and false negative rates (unwanted transitions from the analyte-present region). In Appendix B.3, we show the auxiliary functions to get Proposition 1 from Theorem 2:

#### Proposition 1

(Boundary Distribution Bounds). *Let* **X** *be a bounded birth-death CTMC on X* = 0, 1, …, *P*. *Consider two cut-off processes:*

- 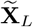 *on V*_*L*_ ∪ *{a*_*L*_*} with a*_*L*_ > *V*_*L*_ ⊂ *χ*, *decay parameter λ*_*L*_, *and* 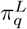
- 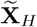 *on V*_*H*_ ∪ *{a*_*H*_ *} with a*_*H*_ < *V*_*H*_ ⊂ *χ*, *decay parameter λ*_*H*_, *and* 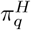

*For boundary initial distributions d*_0_ *(with probability* 1 *of starting at state 0) and d*_*P*_ *(with probability* 1 *of starting at state P):*

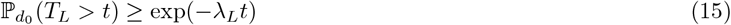

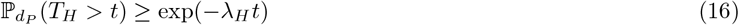

*where T*_*L*_ *and T*_*H*_ *are the exit times from V*_*L*_ *and V*_*H*_ *respectively*.

### 2.3 Applications

In this section, we apply our framework to a Bi-Stab;e Toggle Switch, other applications such as biosensors and non-bi-stable GRNs can be found in Appendix A.

#### 2.3.1 BCRN Bi-Stable Toggle Switch System

A BCRN bi-stable toggle switch is a type of Biochemical Reaction Network characterized by the presence of two distinct and stable equilibrium states [19, 17]. The name “switch” derives from its potential to transition between those two stable states either (typically intentional) with an external perturbation or (typically unintentional) a purely stochastic transition. As a model, we may use two BCRN species of proteins, S_1_ and S_2_, and their respective producers, P_1_ and P_2_. Species S_1_ represses the production of S_2_ and vice-versa, creating the two stable states: either a high S_1_ low S_2_ count or a high S_2_ low S_1_ count. We note their respective counts as *S*_1_, *S*_2_, *P*_1_, *P*_2_, and *n*_*h*_ as the Hill coefficient (cooperative coefficient).

In this section, we employ our framework to assess the probability that the toggle switch moves from the vicinity of one stable state to the other due to stochastic fluctuations (stochastic switching). Below, we state the model of the BCRN toggle switch used for this analysis:

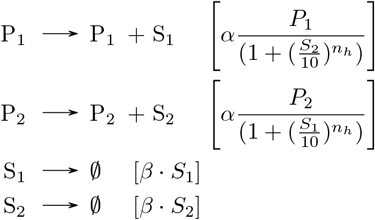

Although it often appears in Ordinary Differential Equation (ODE) form, the toggle switch model presented here follows the widely-adopted model using Hill functions for mutual repression dynamics [19, 11, 35, 45, 29, 32].

As we consider the values of P_1_ and P_2_ to be constants, the states are solely determined by the counts *S*_1_ and *S*_2_ with a state space *χ* = ℕ^2^ Let *S*_1_ = *x* ∈ ℕ and *S*_2_ = *y* ∈ ℕ. The CTMC derived from this BCRN is:

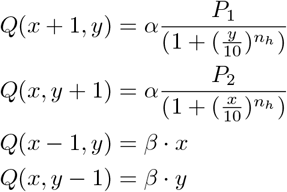

For our analysis, we consider the symmetric case where *P*_1_ = *P*_2_ = *P*, representing equal producer counts for both components of the toggle switch.

We chose *α* = 0.05 s ^−1^ and *β* = 0.01 s ^−1^ as parameters to fall within the range of values used to describe a strong promoter (i.e., the gene regulator) while the degradation rate *β* corresponds to protein half-lives of about 1 minute which is usually not typical for bacteria [1, 33]. The idea behind this excessive parametrization is to provide a stress test where stochastic fluctuations are maximized. Most GRNs would be modeled with weaker promoters and a slower degradation rate, which would imply a lower variance and less likely to have an unwanted transition due to stochastic fluctuations.

The steady-state solution for the deterministic ODE system version described by 17 predicts that two stable equilibrium points exist. The stable equilibrium points occur at 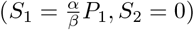 and the other at 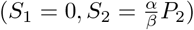.

To predict the QSDs with Theorem 1, we define the set *V* around one of the stable states. For reasons of symmetry, we consider only the state with higher *S*_1_ than *S*_2_ by setting 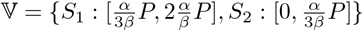. This choice ensures that transitions where 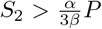 indicate the system has likely switched from the high *S*_1_ to the high *S*_2_ states. The initial condition is set to the deterministic steady state 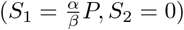 to examine stability under stochastic fluctuations.

We study how both different gene copy numbers {2, 6, 10, 20, 40} and Hill coefficients *n*_*h*_ ∈ {2, 3, 4} determine the susceptibility of the toggle switch to the random stochastic transitions between states. Since 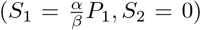, the counts *S*_1_ and *S*_2_ in the steady state are equal to 10, 30, 50, 100, 200 which is within the same range of a reasonable protein count inside a cell. In Table 1, we present these results.

**Table 1:** Toggle Switch Robustness Analysis. Transition probabilities are calculated as 1 − max(0, *e* ^−λ*t*^ −*A*) for different time intervals. The 1*/λ* column represents the characteristic time scale of the QSD.

| $P$ | $n$ | Decay Parameter $s^{-1}$ ($\lambda$) | $1/\lambda$ | $A$ | $\mathbb{P}(T \leq 6\text{h})$ | $\mathbb{P}(T \leq 12\text{h})$ | $\mathbb{P}(T \leq 24\text{h})$ |
| --- | --- | --- | --- | --- | --- | --- | --- |
| 2 | 2 | $1.74 \cdot 10^{-2}$ | 57.4 s | 0.14 | 1 | 1 | 1 |
| 6 | 2 | $1.93 \cdot 10^{-3}$ | 8.7 min | $4.62 \cdot 10^{-2}$ | 1 | 1 | 1 |
| 10 | 2 | $4.33 \cdot 10^{-5}$ | 6.4 h | $1.68 \cdot 10^{-2}$ | 0.62 | 0.86 | 0.99 |
| 20 | 2 | $4.4 \cdot 10^{-10}$ | 72.1 years | $3.99 \cdot 10^{-3}$ | $4 \cdot 10^{-3}$ | $4.01 \cdot 10^{-3}$ | $4.02 \cdot 10^{-3}$ |
| 40 | 2 | 0 | ∞ | $1.45 \cdot 10^{-3}$ | $1.45 \cdot 10^{-3}$ | $1.45 \cdot 10^{-3}$ | $1.45 \cdot 10^{-3}$ |
| 2 | 3 | $1.41 \cdot 10^{-2}$ | 1.2 min | 0.14 | 1 | 1 | 1 |
| 6 | 3 | $1.03 \cdot 10^{-4}$ | 2.7 h | $1.2 \cdot 10^{-2}$ | 0.91 | 1 | 1 |
| 10 | 3 | $3.29 \cdot 10^{-9}$ | 9.6 years | $5.8 \cdot 10^{-3}$ | $5.87 \cdot 10^{-3}$ | $5.94 \cdot 10^{-3}$ | $6.08 \cdot 10^{-3}$ |
| 20 | 3 | 0 | 31989283.5 years | $2.17 \cdot 10^{-3}$ | $2.17 \cdot 10^{-3}$ | $2.17 \cdot 10^{-3}$ | $2.17 \cdot 10^{-3}$ |
| 40 | 3 | 0 | ∞ | $7.55 \cdot 10^{-4}$ | 0 | 0 | 0 |
| 2 | 4 | $1.18 \cdot 10^{-2}$ | 1.4 min | 0.14 | 1 | 1 | 1 |
| 6 | 4 | $8.33 \cdot 10^{-6}$ | 1.4 days | $1.16 \cdot 10^{-2}$ | 0.18 | 0.31 | 0.52 |
| 10 | 4 | $1.52 \cdot 10^{-9}$ | 20.8 years | $4.89 \cdot 10^{-3}$ | $4.92 \cdot 10^{-3}$ | $4.95 \cdot 10^{-3}$ | $5.02 \cdot 10^{-3}$ |
| 20 | 4 | 0 | ∞ | $1.53 \cdot 10^{-3}$ | $1.53 \cdot 10^{-3}$ | $1.53 \cdot 10^{-3}$ | $1.53 \cdot 10^{-3}$ |
| 40 | 4 | 0 | ∞ | $6.26 \cdot 10^{-4}$ | 0 | 0 | 0 |

The results in Table 1 offer design recommendations for engineering GRNs. For all Hill coefficients and gene copy numbers *P* = 20, 40, the probability of the system leaving a stable state due to stochastic fluctuations alone is virtually nonexistent. Notably, higher *n*_*h*_ values allow fewer gene copies for stability. For lower gene copies, as this is a probability upper bound our method’s high bounds don’t rule out stochastic switching.

Finally, for illustrative purposes with more detailed calculations, we demonstrate the QSD using *α* = 0.001 s ^−1^, *β* = 0.001 s ^−1^, *P* = 30 genes, and 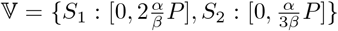. These parameters, representing weaker promoters with balanced production and degradation rates, are more suitable for QSD visualization. First, the resulting steady-state *S*_1_ counts (30 molecules) create a system where stochastic fluctuations are significant relative to the mean, making quasi-stationary behavior more pronounced. Second, the smaller state space facilitates complete numerical calculation of the QSD while remaining biologically relevant. Finally, the system’s tendency to remain longer within *V* before exiting allows the QSD structure to fully develop, providing a clearer illustration of the distribution around the two stable states.

For this choice of *V*, the decay parameter is 8.34 *·* 10 ^−8^ and a bound constant *A* of 0.005. In Figure 2, we calculate two QSDs one for *S*_1_’s states probability distribution and the other for *S*_2_’s. The predicted QSDs both yield a total variation distance of 3.6 *·* 10 ^−3^ with the original processes’ time evolution.

**Figure 2.**
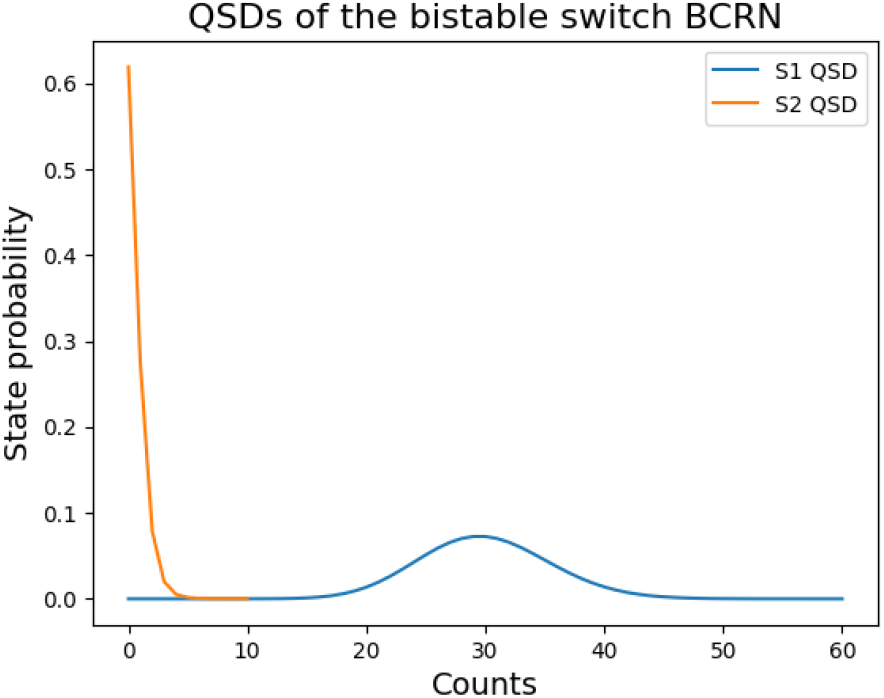
Quasi-steady State Distributions: This figure presents the quasi-steady state probability distributions predicted for the toggle switch with *α* = 0.001 s ^−1^, *β* = 0.001 s ^−1^, *P* = 30, and *V*

This toggle switch analysis demonstrates our framework’s ability to provide design principles for GRNs. Our results match the current literature and indicate that GRNs with higher gene copy numbers are more resistant to stochastic switching [19, 30, 31, 41]. It is important to consider that the number of steady-state protein concentrations (or basal level expression) is also a highly important factor to consider, as it directly gives the number of states that separate the vicinity of one stable state from the other. A gene with a lower basal level expression would not perform as well even with more than 20 copies, as is indicated by CTMC theory [44] and how increasing the number of steady state proteins reduces the probability of stochastic switching. Finally, it is important to emphasize that our analysis was limited to a single stress-test model designed to maximize stochastic fluctuations through excessive parametrization. As GRNs would be modelled with more moderate parameters, implying reduced susceptibility to unwanted transitions. For circuit-specific design recommendations, we recommend using our computational framework with parameters appropriate to individual systems rather than relying on these stress-test bounds.

## 3 Methods

### 3.1 Proof Strategy for Theorem 1

The strategy uses the law of total probability:

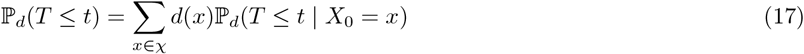

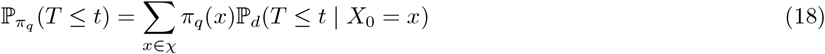

Let *K*_*dif*_ be defined as the following:

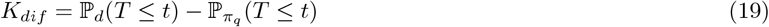

It is verified in the appendix that any constant greater and equal to *K*_*dif*_ for all times *t* satisfies Inequality (11). The constant *A* calculated at the end of the algorithm is equivalent to using the law of total probability and bounding *K*_*dif*_ with 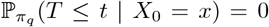 and ℙ_*d*_(*T* ≤ *t* |*X*_0_ = *x*) = 1 for all *x* ∈ *χ* and summing the results with Σ_*x* ∈ *X*_ *d*(*x*). Finally, one can show for the set *A*_*x*_ of states that are directly connected to *x* (i.e. all *z* ∈ *χ* such that *Q*(*x, z*) > 0) the following hold for the jump chain *J* calculated from *Q*:

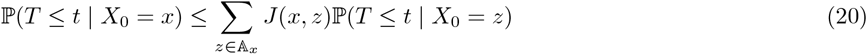

Finally, Equation (20) is used to prove that *D* decreases at each iteration in Algorithm 1, and show that after each step of the algorithm, the constant *A* calculated remains an upper bound to *K*_*dif*_ and the inequality still holds. The complete detailed proof can be found in Appendix B.2.

### 3.2 Proof Strategy for Theorem 2

The strategy lies on proving that the exit time *T* for process 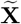 given two states *x* ∈ *χ* and *y* ∈ *χ* with *x* > *y* follows:

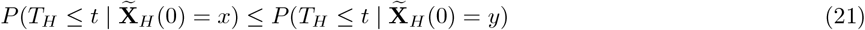

Then, it is verifiable through the law of total probability that the initial distribution *d*_*B*_, where *B* denotes the boundary state furthest from the absorbing state (i.e., state *P* for process 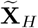 with *a*_*H*_ < *P*, or state 0 for process 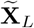 with *a*_*L*_ > 0), has the lowest exit time probability by time *t* compared to any other initial distributions, including the quasi-stationary distribution 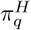, which yields the theorem’s main result. The proof is symmetric, so it also holds for process 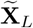. The complete detailed proof can be found in Appendix B.3.

### 3.3 Implementation

For the implementation of Algorithm 1 and QSD estimation we modified the MobsPy compiler [10]. The MobsPy compiler creates an SBML file [25] through a generated Python dictionary with all reactions, while our repository creates a new rate dictionary containing the rate function and the state transition for each reaction. Afterward, we loop through the BCRN states and use the rate dictionary for each state to calculate all transition rates between states. To address the state space explosion problem inherent in multi-species BCRNs, we construct the generator matrix 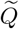 of the cut-off process in *V* directly as a sparse matrix. This approach avoids the memory overhead of dense matrix construction and enables analysis of systems with hundreds of thousands of states.

The script then performs a connectivity check using the sparse 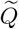 starting from the state with the smallest species counts *s*_*c*_ in set *V*. Connectivity is verified by checking if every state is reachable from *s*_*c*_, then by checking if every state can reach *s*_*c*_ through the transpose of matrix 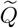. If the connectivity check is passed, the QSD is calculated by constructing the conditional generator matrix 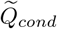 with the rates to the absorbing state removed. The matrix diagonal is set to ensure proper row sums (i.e., 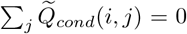 for all *i*), maintaining the fundamental property of generator matrices.

The decay parameter *λ* is estimated directly from the eigenvalues of 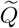 using sparse eigenvalue solvers. Specifically, *λ* is the negative of the largest real eigenvalue of 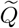, corresponding to the principal eigenvalue of the quasi-stationary distribution. The QSD *π*_*q*_ is obtained as the normalized left eigenvector corresponding to this principal eigenvalue. The sparse eigenvalue-based approach provides numerical stability and computational efficiency compared to matrix exponential methods, particularly for large sparse systems where direct exponentiation is impossible.

The sparse matrix implementation, combined with efficient eigenvalue computation, enables the analysis of toggle switch systems with state spaces exceeding 600, 000 states, which would be intractable using dense matrix methods.

## Supporting information

Supplemental Results/Mathematical Proofs

## Acknowledgements

The work was supported by the French National Research Agency (ANR) projects DREAMY (ANR-21-CE48-0003), COSTXPRESS (ANR-23-CE45-0013), the SAIF project, funded by the “France 2030” government investment plan managed by ANR, under the reference ANR-23-PEIA-0006 (M.F., T.N.) and the National Institute of Mental Health (K99 MH130894; R00 MH130894) granted to Dr. Stephanie Noble (F.C.).

