## Supplemental Results/Mathematical Proofs for "A Quasi-Stationary Distribution Bound for Fault Analysis in Gene Regulatory Networks"

### A Extended Results

#### A.1 An Allee-Based Algorithm For Rare Event Detection

In previous work [9], we proposed a BCRN modeled with a deterministic ordinary differential equation (ODE) system that coordinates cells to detect an analyte of interest called the Allee-based algorithm. In this BCRN, cells transition between a logical low L to a logical high H in response to contact with the analyte. Cells from the high H state can convert cells from low L to high H, and as their number increases, so does this rate of conversion. A cell can switch back from the high H state to the low L to prevent the accumulation of incorrect detection events. If the analyte concentration is low, two stable states are present: one with a lower count of H cells and another with a higher count. Conversely, if the analyte concentration is high enough, only the higher H stable state is present, and the ODE system converges to this state. The higher H stable state also functions as a memory element; if the analyte concentration starts high and then decreases, the ODE system will still converge to the higher H stable state. Therefore, the stable states indicate the presence or absence of the analyte, with the lower H state indicating absence and the higher H state indicating presence.

The Allee-based algorithm comprises three reactions: (i) Reaction (Detect): A cell in state L changes to state H upon contact with the analyte with a rate $\sigma_A \in \mathbb{R}_+$. (ii) Reaction (Hold): A cell in state L switches to state H with a rate that depends on the number of H cells according to a Hill function with parameters $\kappa \in \mathbb{R}_{>0}$, $K \in \mathbb{R}_{>0}$, and $n \in \mathbb{R}$ with $n > 1$. The Hill function models the reaction triggered by a QS molecule, which is secreted by cells in the H state; see, e.g., [14] for a Hill-function model of a QS circuit. (iii) Reaction (Reset): A cell in state H switches back to state L with a certain *reset rate* $\rho \in \mathbb{R}_{>0}$. Intuitively, this hinders the incorrect detection events in the system. Finally, note as $P \in \mathbb{N}$ the number of cells in the population.

$$\mathrm{L} + \mathrm{A} \longrightarrow \mathrm{H} + \mathrm{A} \quad [\sigma_A \cdot A \cdot L] \tag{Detect}$$

$$\mathrm{L} + \mathrm{H} \longrightarrow 2 \cdot \mathrm{H} \quad \left[\kappa \cdot \frac{H^n}{H^n + K^n} \cdot L\right] \tag{Hold}$$

$$\mathrm{H} \longrightarrow \mathrm{L} \quad [\rho \cdot H] \tag{Reset}$$

(22)

For simplification, we consider a constant analyte concentration and rewrite reaction (Detect) as

$$\mathrm{L} \longrightarrow \mathrm{H} \quad [\sigma \cdot L] \tag{Set}$$

Let $x \in \chi$ with $0 \leq x \leq P$. As the population value $P$ is constant when replication and death are not considered, one has $L = (P - H)$. From the Allee-based algorithm BCRN, the respective CMTP follows:

$$Q(x, x+1) = \sigma \cdot (P - H) + \kappa \cdot \frac{H^n}{H^n + K^n} \cdot (P - H)$$
$$Q(x, x-1) = \rho \cdot H$$

Here, we model the BCRN stochastically with a CMTP model instead of the deterministic ordinary differential equation (ODE) system originally used to describe its dynamics [9]. Typically, ODE models are suitable when BCRN species have much higher counts, as suggested by Kurtz's results on the convergence of stochastic processes to their deterministic ODE counterparts [28]. Further, ODE models typically use the concentration of the BCRN species, while the CTMCs use the counts. Thus, parameters in units of concentration (cells per volume) are multiplied by the volume to convert to the corresponding parameter in terms of counts (cells).

Parameters $H, n, \rho, k$ remain unchanged as they do not depend on cell counts or have concentration units. Refer to Table 2 for their values. The parameter $K$ represents the soft threshold concentration of cells required to trigger the conversion from the low state to the high state by other cells in the high state (Reaction (Hold)) in the deterministic ODE model. As it is a concentration, we can convert $K$ to cell count by multiplying the volume in which cells are contained. We confine the cells to a volume in which $K$ is between 30% and 50% of the number of cells in the population $P$. On the one hand, $K$ cannot be too low to prevent a false positive due to erroneous transitions from L to H without analyte detection. On the other hand, $K$ must not be so high as to stop a true positive when the analyte is present.

We use the QSD probability bound from Theorem 1 to estimate false positive and false negative rates and use this to estimate $P$ to calculate the necessary volume according to Equation (23):

$$K = f \cdot P \tag{23}$$

$$v = \frac{K}{K'} \tag{24}$$

where $K'$ is the concentration-based value of $K$ estimated in our previous work [9] and $f$ is the fraction of the population used for $K$.

| Parameter | Value | Source |
|---|---|---|
| $\kappa$ | $35\text{h}^{-1}$ | ([14]) |
| $\rho$ | $14\text{h}^{-1}$ | ([14]) |
| $n$ | 4 | ([14]) |
| $\sigma$ | $0.5\text{h}^{-1}$ | ([9]) |
| $K$ | 80 cells | estimated |
| $P$ | 200 cells | estimated |
| $v$ | $1 \cdot 10^{-6}\text{mL}$ | estimated |

Table 2: **Reaction and population parameters used in simulations.**

In our previous work [9], we estimated $\sigma = 0.5\text{h}^{-1}$ to be a baseline error rate that causes the conversion from L to H erroneously without a reaction with the analyte. Under these circumstances, we desire the algorithm to yield the analyte-absent output.

To use Proposition 1, we select $\mathbb{V}_L$ that will encompass the desired states that indicate the absent analyte output using the fact that this BCRN deterministic model possesses three equilibrium points. From our previous results [9], if $H(0) < p_I < P$, where $p_I$ is an unstable equilibrium point between the two stable equilibrium points, the ODE BCRN converges to the lower H stable state and if $H(0) > p_I$, it converges to the higher H. For a visual example, refer to Figure 5 c. Based on these equilibrium points, we propose $\mathbb{V}_L = \{0, 1, 2, ...., \lceil p_I \rceil\}$. The parameters from Table 2 yield $\lceil p_I \rceil = 48$ cells. To define the initial distribution, let $d_0$ be as follows:

$$\mathbb{P}_{d_0}(H(0) = 0) = 1 \tag{25}$$

Additionally, let $\lambda_L$ be the decay parameter from the generator matrix of the cut-process in $\mathbb{V}_L$.

We consider a false positive if there was ever a time during the execution of the algorithm where the cell population is in a state outside of $\mathbb{V}_L$ with $d_0$ as an initial distribution. Therefore, the false positive probability is given by $\mathbb{P}_{d_0}(T_L \leq t_e) = 1 - \mathbb{P}_{d_0}(T_L > t_e)$ where $t_e \in \mathbb{R}$ is the execution time of the algorithm and $T_L$ is the exit time from $\mathbb{V}_L$ (as defined in Proposition 1). We take as $t_e = 24\text{h}$ as it is a reasonable time for an envisaged practical application. Finally, refer to Table 3 for the decay parameter and $\mathbb{P}_{\pi_q^L}(T_L \leq 24\text{h})$ which is the upper bound on the false positive probability from Proposition 1.

After the system reaches the vicinity of the higher H stable state, it must remain to yield a positive output. Otherwise, it would result in a false negative. To estimate the false negative rate with Proposition 1, we define $\mathbb{V}_H$ to encompass the states that yield a positive output using the remaining states from $H = \lceil p_I \rceil + 1$ to $H = P$.

We consider a false negative if there was ever a time during the execution of the algorithm where the cell population is in a state outside of $\mathbb{V}_H$ with $d_P$ as an initial distribution, where $d_P$ is the distribution given by:

$$\mathbb{P}_{d_P}(H(0) = P) = 1 \tag{26}$$

Additionally, let $\lambda_H$ be the decay parameter from the generator matrix of the cut-process in $\mathbb{V}_H$. Finally, we repeat the calculations performed for Table 3 to create Table 4.

Values smaller than $10^{-15}$ in Tables 3 and 4 are treated as numerically zero due to machine precision limitations in eigenvalue calculations for generator matrices. These ultra-small decay parameters indicate that the corresponding

| | $\lambda_L$ h$^{-1}$ | | $\mathbb{P}_{\pi_q^L}(T_L \leq 24$ h$)$ | |
|---|---|---|---|---|
| P | $K = 0.3 \cdot P$ | $K = 0.5 \cdot P$ | $K = 0.3 \cdot P$ | $K = 0.5 \cdot P$ |
| 20 | 0.84 | $1.42 \cdot 10^{-2}$ | 1.0 | 0.29 |
| 60 | 0.16 | $2.76 \cdot 10^{-6}$ | 0.98 | $6.63 \cdot 10^{-5}$ |
| 100 | $3.02 \cdot 10^{-2}$ | $4.79 \cdot 10^{-10}$ | 0.52 | $1.15 \cdot 10^{-8}$ |
| 140 | $5.75 \cdot 10^{-3}$ | $6.51 \cdot 10^{-14}$ | 0.13 | $1.56 \cdot 10^{-12}$ |
| 200 | $4.66 \cdot 10^{-4}$ | 0 | $1.11 \cdot 10^{-2}$ | 0 |
| 400 | $1.01 \cdot 10^{-7}$ | 0 | $2.43 \cdot 10^{-6}$ | 0 |
| 750 | $2.08 \cdot 10^{-14}$ | $1.37 \cdot 10^{-14}$ | $4.99 \cdot 10^{-13}$ | $3.3 \cdot 10^{-13}$ |

Table 3: **Estimated false positive rates for the Allee-Based Algorithm**: Upper bounds of the false negative probability $\mathbb{P}_{\pi_q^L}(T_L < 24\text{h})$, which is the probability that the cellular population will leave the set of states $\mathbb{V}_L$ that represent a negative output, for different values of $K$ and $P$.

| | $\lambda_H$h$^{-1}$ | | $\mathbb{P}_{\pi_q^H}(T_H \leq 24\text{h})$ | |
|---|---|---|---|---|
| P | $K = 0.3 \cdot P$ | $K = 0.5 \cdot P$ | $K = 0.3 \cdot P$ | $K = 0.5 \cdot P$ |
| 20 | $6.78 \cdot 10^{-3}$ | 4.36 | 0.15 | 1.0 |
| 60 | $7.42 \cdot 10^{-10}$ | 0.42 | $1.78 \cdot 10^{-8}$ | 1.0 |
| 100 | 0 | $5.73 \cdot 10^{-2}$ | 0 | 0.75 |
| 140 | $1.03 \cdot 10^{-13}$ | $8.91 \cdot 10^{-3}$ | $2.47 \cdot 10^{-12}$ | 0.19 |
| 200 | $9.84 \cdot 10^{-14}$ | $4.4 \cdot 10^{-4}$ | $2.36 \cdot 10^{-12}$ | $1.05 \cdot 10^{-2}$ |
| 400 | $\leq 1 \cdot 10^{-15}$ | $2.29 \cdot 10^{-8}$ | 0 | $5.49 \cdot 10^{-7}$ |
| 750 | $\leq 1 \cdot 10^{-15}$ | 0 | 0 | 0 |

Table 4: **Estimated false negative rates for the Allee-Based Algorithm**: Upper bounds for the false negative probability $\mathbb{P}_{\pi_q^H}(T_H \leq 24\text{h})$, which is the probability that the cellular population will leave the set of states $\mathbb{V}_H$ that represent a positive output, for different values of $K$ and $P$.

transitions occur on timescales of millions of years or longer, making them practically irrelevant for biological applications. The mathematical framework correctly identifies these systems as essentially stable under realistic experimental conditions.

From Tables 3 and 4, one can see $K = 0.3 \cdot P$ better avoids false positives, while $K = 0.5 \cdot P$ better avoids false negatives, with higher population values contributing for avoiding both. Therefore, we choose a population value of $P = 200$ cells with $K = 0.4 \cdot P = 80$ cells, as it assures both estimated bounds for the false positive and false negative probabilities: $\mathbb{P}_{d_0}(T_L \leq 24\text{h}) \approx 0$ and $\mathbb{P}_{d_P}(T_H \leq 24\text{h}) \approx 0$. This results in a volume of $1 \cdot 10^{-6}\text{mL}^3$ from Equation (23) and the parameters are presented in Table 2.

For the ODE model (22), if the analyte concentration increases, the stable equilibrium point for the lower H state disappears, and the system converges to the higher H state [9] (see Figure 5). The lowest rate where there are no two stable points is the intended detection rate for the algorithm. This rate is found by a binary search over lower rates, which have two stable equilibrium points, and higher rates, which have only one. We find a rate of $\sigma = 1.8\text{h}^{-1}$ with the parameters from Table 2. Finally, the true positive rate comes from the numerical solution of the Chemical Master Equation for the probability of finding the BCRN in the vicinity of the higher H state $\mathbb{V}_H$ with rates higher than $\sigma = 1.8\text{h}^{-1}$ after 1h starting from the initial distribution $d_0$.

We now compare both QSDs from the cut-process in $\mathbb{V}_L$ and $\mathbb{V}_H$ which we respectively note as $\pi_q^L$ and $\pi_q^H$, with the time evolution of two state probability distributions of the original BCRN $d_{t0}$ and $d_{tP}$. The distribution $d_{t0}$ has $d_0$ as an initial distribution and $d_{tP}$ has $d_P$. The time evolution is given by solving the Chemical Master Equation numerically. It evaluates how accurately the cut-off processes can predict the behavior of the BCRN. Refer to Figure 3 for the comparison.

Figures 3 a, b, c, and d, show that the BCRN approaches $\pi_q^L$ when the initial distribution is $d_0$ and $\pi_q^H$ when its $d_P$. In both cases, convergence is not observed. The calculated decay parameter for the cut-process in $\mathbb{V}_L$ is $2.78 \cdot 10^{-10}$ and

| Set Rate $\sigma$ h$^{-1}$ | Higher H State Vicinity $\mathbb{V}_H$ Probability |
|---|---|
| 1.8 | 0.70 |
| 2 | 0.93 |
| 3 | 1.0 |
| 5 | 1.0 |

Table 5: **Estimated True Positive Rate for The Allee-Based Algorithm.** Probability of finding the cell population in a state in $\mathbb{V}_H = \{49, 50, 51, ...200\}$ given a rate that indicates the presence of an analyte after one hour has passed and the system started with distribution $d_0$.

$1.11 \cdot 10^{-16}$ for $\mathbb{V}_H$. When bound with Proposition 1, it indicates that both $\mathbb{P}(T_L \leq t)$ and $\mathbb{P}(T_H \leq t)$ are lower than 0.0003 for the simulated time frames. As predicted by Equation (8), the low values of $\mathbb{P}(T_H \leq t)$ and $\mathbb{P}(T_L \leq t)$ allow the original process to approach the predicted QSD by the cut-off process. The slight difference between the QSD and the time evolution lies in a non-zero probability of the BCRN being in states not in $\mathbb{V}_L$ or $\mathbb{V}_H$ for the lower H and higher H cases, respectively. Moreover, only after $10^5$h one can observe a shift in the original distribution from states in $\mathbb{V}_L$ to states in $\mathbb{V}_H$, a time scale far beyond the intended operation of the algorithm.

A formula to solve for the steady-state distribution of this BCRN is provided by Karlin and Taylor [24] with the exact analytical solution in Figure 4. We required times up to $t = 10^{13}$h when attempting to simulate the stationary distribution by solving the matrix exponential of the BCRN using Python's `scipy.linalg.expm`. Times up to $t = 10^{15}$h yield more numerical errors with distributions with a probability sum greater than one. This indicates that some BCRNs with quasi-stationary distributions that converge to stationary distributions present challenges when simulating their stationary distribution numerically due to the extensive timescales involved.

In Figure 5 a and b, we show the effects of the initial distribution on the probability of the BCRN being in each state. Distributions around the unstable equilibrium point divide their total probability into $\mathbb{V}_L$ and $\mathbb{V}_H$ with shapes resembling the ones of $\pi_q^L$ and $\pi_q^H$, The distribution with the probability division remains for a time scale similar to the duration of the quasi-stationary distribution before the convergence to the steady-state distribution, showing that more than two distributions remain almost stationary before converging to a steady state.

Finally, we use Algorithm 1 to estimate the constant $A$ for different initial distributions. In Table 6, we compare the obtained $A$ from the execution of Algorithm 1 and the total variation distance between the time evolution of the state distribution after 24h and the QSD $\pi_q^L$. One can notice that higher $A$ values correlate with a higher total variation distance between the QSD $\pi_q^L$ as predicted by Equation (8). The similar $A$ value for $H$ of 5, 10, and 20 is due to the algorithm stopping after tolerance $\epsilon$ is satisfied.

| Initial Distribution | $A$ | Total Variation Distance: $\pi_q^L$ and Time Evolution for 24h |
|---|---|---|
| $H = 5$ | $9.97 \cdot 10^{-6}$ | $1.8 \cdot 10^{-4}$ |
| $H = 10$ | $9.99 \cdot 10^{-6}$ | $1.8 \cdot 10^{-4}$ |
| $H = 20$ | $9.98 \cdot 10^{-6}$ | $1.8 \cdot 10^{-4}$ |
| $H = 40$ | $10.0 \cdot 10^{-1}$ | $6.1 \cdot 10^{-2}$ |
| $H = 47$ | 1.0 | $4.5 \cdot 10^{-1}$ |

Table 6: **The A Constant vs. Initial Distribution**. Values of A for different initial distributions. The distributions have probability one being in the state indicated by the equality with $H$. We further calculate the total variation distance between the time evolution of the state probability distribution after 24h and the cut-off process QSD ($\pi_q^L$) for different initial distributions.

In this subsection, we demonstrated the efficiency of the cut-off process when analyzing BCRNs with multiple stable states using the Allee-based algorithm. The quasi-stationary distribution of the cut-off process proved to be an excellent predictor of BCRN behavior for specific initial distributions. Additionally, the decay parameter and Theorem 2 provided an efficient probability bound on false positives and false negatives, facilitating the parametrization of the Allee-based algorithm. Moreover, we highlighted the significance of considering the initial distribution by showing that distributions

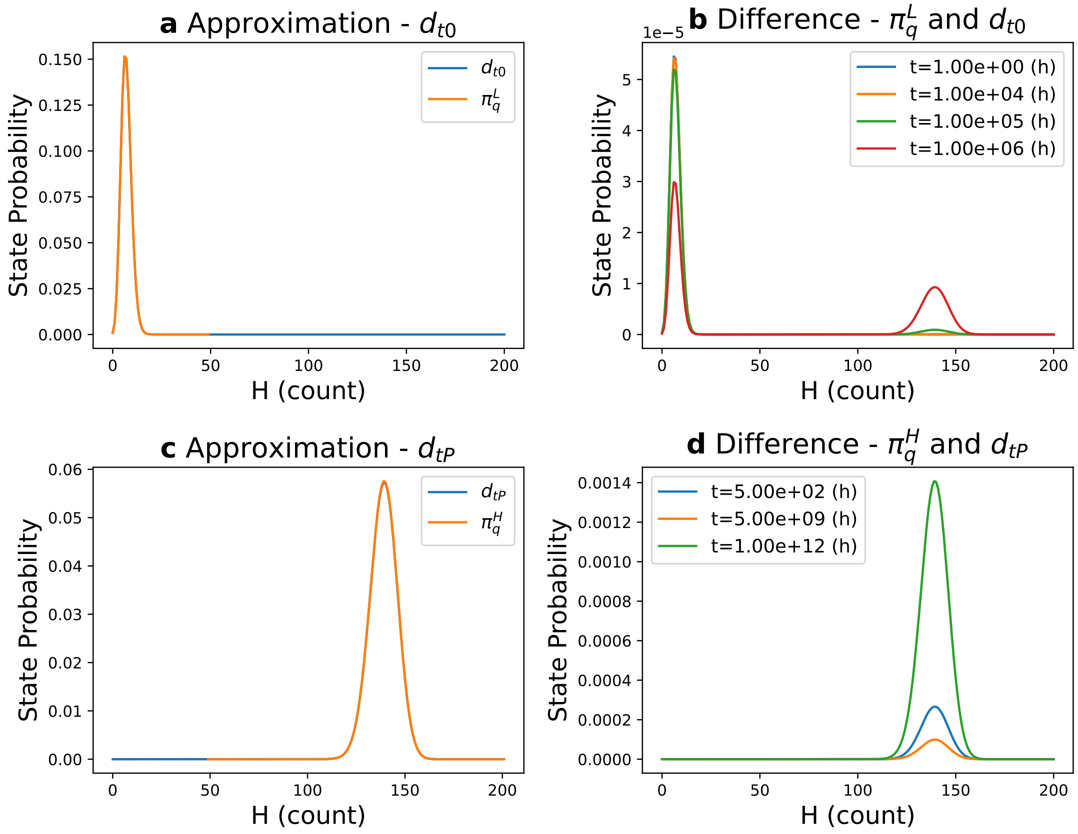


Figure 3: **Comparison with Quasi-Stationary Distributions.** **(a) Comparison between $d_{t0}$ and $\pi_q^L$:** Time evolution of the state probability distribution of the original BCRN $d_{t0}$ with $d_0$ as an initial distribution after $10^3$h with the QSD of the cut-off process in $\mathbb{V}_L$ ($\pi_q^L$). **(b) Difference between $\pi_q^L$ and $d_{t0}$:** Absolute difference between the time evolution of the state probability distribution of the original BCRN $d_{t0}$ with $d_0$ as an initial distribution and QSD of the cut-off process in $\mathbb{V}_L$ for the times of 1h, $10^4$h, $10^5$h, and $10^6$h. **(c) Comparison between $d_{tP}$ and $\pi_q^H$:** Time evolution of the state probability distribution of the original BCRN $d_{tP}$ with $d_0$ as an initial distribution after $10^3$h with the QSD of the cut-off process in $\mathbb{V}_H$ ($\pi_q^H$). **(d) Difference between $\pi_q^H$ and $d_{tP}$:** Absolute difference between the time evolution of the state probability distribution of the original BCRN $d_{tP}$ with $d_P$ as an initial distribution and QSD of the cut-off process in $\mathbb{V}_H$ for the times of 500h, $5 \cdot 10^9$h, and $1 \cdot 10^{12}$h.

concentrated closer to $p_I$ yield quasi-stationary distributions, which remain almost stationary and distributed between the two stable states. Algorithm 1 indicated higher $A$ values for these distributions.

##### A.2 Controlling Spatiotemporal Pattern Formation in a Concentration Gradient with a Synthetic Toggle Switch

Barbier et al. [4] proposed a BCRN to generate spatial color patterns based on the concentration of two BCRN species, AHL and IPTG. It functions with the interplay of two other chemical species, LacI and TetR, which mutually repress each other. Unlike the previous bistable examples, this system exhibits either monostability or bistability depending on the local concentrations of IPTG and AHL. LacI, when chemically bound with IPTG, represses the production of TetR, which results in IPTG controlling the strength of the repression of TetR. Moreover, AHL triggers TetR production.

Below, we provide the MobsPy [10] implementation, as the complexity of GRN, with its multiple repression mecha-

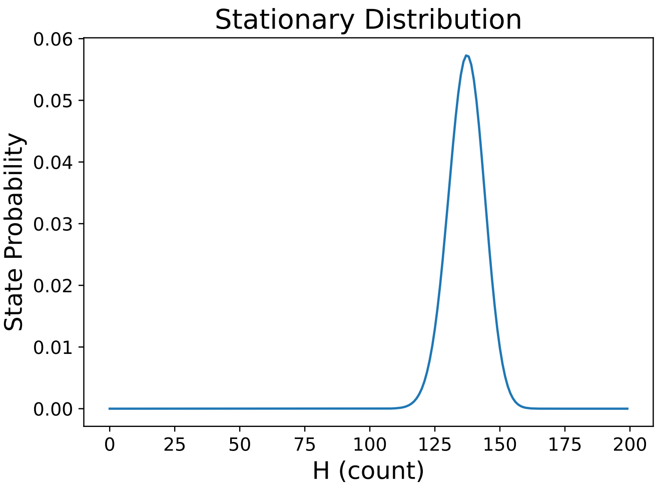


Figure 4: **Steady-State Distribution of the Allee-Based Algorithm.** Analytically solved steady-state distribution of the Allee-Based Algorithm. On the x-axis, there is the number of H cells, and on the y-axis, the probability for the system to have that number of cells in the H state.

nisms and conditional rate functions, requires some synthesis as opposed to the simpler models used so far.

We scaled the original parameters from [4] by multiplying the repression thresholds (Kt, Kl) by 5 to strengthen bistability and the AHL sensitivity (Ka) by 0.05 to reduce switching frequency, transforming transition timescales from $10^8$ arbitrary time units to tractable simulation ranges while checking that it preserved the multi-stability properties of the original circuit. These changes were performed to make sure the simulations would perform better for a stochastic model. Given this scaling potential, we leave the model in arbitrary units to avoid misinterpretation of the results.

```
by, bx, Ki, Kt, Kl, Ka, dy, dx, nt, na, nl = 43.8, 36.2, 2.22, 27.4*5, \
                                            4.17*5, 0.133*0.05, 1, 1, 2.29, 1.61, 2.17

LacI, TetR, IPTG, AHL = BaseSpecies()

# LacI repression and TetR decay
TetR >> TetR + LacI [lambda r: repression_rate(r, None, None, by, Kt, nt, None, None, None)]
TetR >> Zero [dx]

# TetR repression and LacI
rate_f = lambda r, r2, r3: repression_rate(r, r2, r3, bx, Kl, nl, Ki, Ka, na)
LacI + IPTG + AHL >> LacI + IPTG + AHL + TetR [rate_f]
LacI >> Zero [dy]

def repression_rate(r, r2, r3, b, K, n, K_I, K_ahl, n_ahl):
    if r2 is not None:
        r = r/(1 + K_I*r2)
    if r3 is not None:
        f = (K_ahl*r3)**n_ahl/(1 + (K_ahl*r3)**n_ahl)
    return b/(1 + (K*r)**n)*f if r3 is not None else b/(1 + (K*r)**n)

model = set_counts({LacI: 45, TetR: 0, IPTG: 0, AHL: 1})
interval = {LacI: [200, 10], TetR: [0, 8], IPTG: [0, 0], AHL: [1, 1]}
Sim = Simulation(model)
J = Jump_chain_qsd_bound(Sim, interval)
```

The authors concluded there was an irreversible high LacI state for low values of IPTG in their deterministic model. We propose the set $\mathbb{V} = \{\text{LacI} : [10, 200], \text{TetR} : [0, 8], \text{IPTG} : [0, 0], \text{AHL} : [1, 1]\}$ corresponding to the monostable high LacI regime, and analyze the probability of stochastic escape from this region to a bistable one. For this model, the

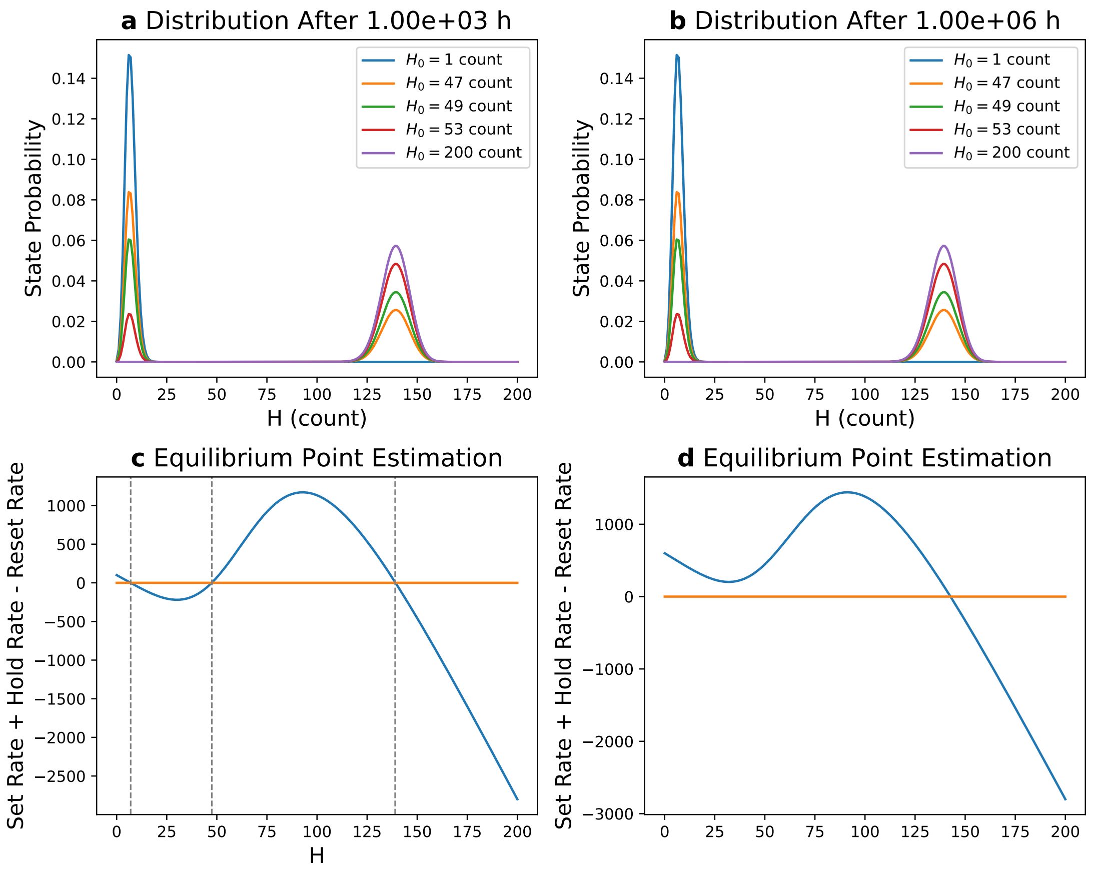


Figure 5: **Time Evolution of State Probability Distributions for Different Initial Values. (a) Time Evolution of the State Probability Distribution After $10^3$h**: Evolution of the probability of the original process being in each state given an initial count $H_0$ after $10^3$h. **(b) Time Evolution of the State Probability Distribution After $1^6$h**: Evolution of the probability of the original process being in each state given an initial count $H_0$ after $10^6$h. There are no noticeable changes between the state probabilities compared to the $10^3$h figure. **(c) Equilibrium Point Visualization**: Plot showing the sum of the Reaction Set rate plus Reaction Hold rate minus Reaction Reset. The zeros in this plot are the equilibrium points. **(d) Loss of Equilibrium Point Visualization**: Plot showing how increasing the Reaction Set rate resulting in two of the equilibrium points disappearing.

calculated decay parameter is $6.35 \cdot 10^{-9}$ with constant $A = 3.3 \cdot 10^{-3}$ for the initial distribution $(\text{LacI} = 45, \text{TetR} = 0, \text{IPTG} = 0, \text{AHL} = 1)$. The probability of the BCRN leaving the set $\mathbb{V}$ cannot be simulated to time $1.34 \cdot 10^8$ (equal to $1/\lambda$) as the ODE solver exhausts its maximum allowed integration steps trying to maintain numerical accuracy across eight orders of magnitude in time. Reducing the step size to address this issue results in each simulation run taking about one hour, making it impractical to simulate 1000 runs for an exit probability estimation. The estimated partial QSD over LacI can be found in Figure 6.

This model demonstrates the application of the QSD bound to study GRNs with multi-stable states, as presented in the literature. The cut-off process successfully predicted the QSD, and the decay parameter and constant $A$ from Algorithm 1 estimated exit times for a state classified as irreversible. Compared to solving the Chemical Master Equation numerically, our method provides significant advantages, as the state space of this BCRN is infinite while the cut-off process is finite. Additionally, the challenge of simulating extensive time scales using the Gillespie Algorithm evidences the importance of less computationally expensive methods for estimating exit times in multi-stable systems.

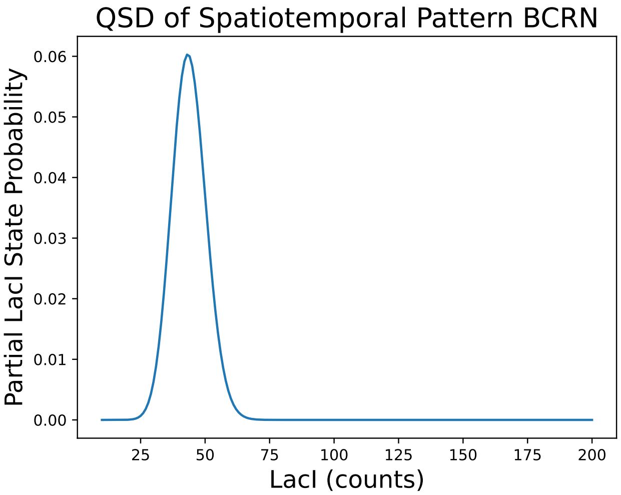


Figure 6: **Partial QSD of LacI**: Partial QSD distribution obtained using the cut-off process $\mathbb{V}$ for the spatiotemporal synthetic toggle switch.

### B Proof of Theorem 1

#### B.1 Proof of Equality Between Exit Times

Let $\mathbf{X}$ be a continuous-time Markov process with a countable sequence of states $\mathbf{X} = (X_0, X_1, X_2, X_3, ...)$ such that $X_i \in \chi$ and a sequence of times $\mathbf{t} = (0, t_0, t_1, t_2, t_3, ...)$ such that $[t_{i-1}, t_i[$ is the interval of time process $\mathbf{X}$ spends in state $X_i$. Let the state space $\chi$ be connected for process $\mathbf{X}$, which means that for every two states $x, y \in \chi$, there exists a path from $x$ to $y$. Let $\mathbf{X}(t)$ be the state of process $\mathbf{X}$ at time $t$. Let $Q$ be the generator matrix of a process $\mathbf{X}$. Let $x, y$ be any two states such that $x, y \in \chi$. The definition of $Q$ follows:

$$\begin{aligned} &Q(x,y) \geq 0 \text{ if } y \neq x \\ &Q(x,x) = -\sum_{y \in \chi} Q(x,y) \end{aligned} \tag{27}$$

Let $d$ be the initial distribution of process $\mathbf{X}$, such that $d(x)$ is the probability that process $\mathbf{X}$ is initially in state $x \in \chi$. From the law of total probability, the probability of finding the process in state $y \in \chi$, given the initial distribution $d$, at state $X_i$:

$$\mathbb{P}_d(X_i = y) = \sum_{x \in \chi} \mathbb{P}(X_i = y \mid X_0 = x) d(x) \tag{28}$$

We now define the connected finite set $\mathbb{V} \subset \chi$ and the singleton $\{a\}$ with $\{a\} \cap \mathbb{V} = \emptyset$. We use the set $\mathbb{V}$ to define a continuous-time Markov process $\widetilde{\mathbf{X}}$ using $\mathbf{X}$. Let the sequence of states $\widetilde{\mathbf{X}} = (X_0, X_1, X_2, X_3, ...)$ be the sequence states of process $\widetilde{\mathbf{X}}$ in the state space $\mathbb{V} \cup \{a\}$, where $\mathbb{V}$ is connected for process $\widetilde{\mathbf{X}}$ and the state $a$ is absorbing. Let the sequence of times $^{\mathbf{x}}\widetilde{\mathbf{t}} = (0, \widetilde{t}_0, \widetilde{t}_1, \widetilde{t}_2, \widetilde{t}_3, ...)$ be the sequence such that $[\widetilde{t}_{i-1}, \widetilde{t}_i[$ is the time process $\widetilde{\mathbf{X}}$ spends in state $\widetilde{X}_i$. Let $\widetilde{\mathbf{X}}(t)$ be the state process of $\widetilde{\mathbf{X}}$ at time $t$. Let $\widetilde{Q}$ be the transition rate matrix of a process $\widetilde{\mathbf{X}}$. Let $x, y \in \mathbb{V}$, $z \in \chi$, and $a \in \chi \setminus \mathbb{V}$.

The definition of $\widetilde{Q}$ follows:

$$\begin{aligned}
\widetilde{Q}(x,y) &= Q(x,y) \quad \text{if } x \neq y \\
\widetilde{Q}(x,a) &= \sum_{w \notin \mathbb{V}} Q(x,w) \\
\widetilde{Q}(a,z) &= 0 \\
\widetilde{Q}(z,z) &= -\sum_{w \in \chi, w \neq z} \widetilde{Q}(z,w)
\end{aligned} \tag{29}$$

We call $\widetilde{\mathbf{X}}$ the cut-off process in $\mathbb{V}$ from $\mathbf{X}$. We omit "from $\mathbf{X}$" when it can be inferred from context.

Let the state space $\chi$ be totally ordered. As the set $\chi$ is completely ordered, we chose an order such that the states $v_1 \in \mathbb{V}$ and $v_2 \notin \mathbb{V}$ must follow:

$$v_1 > v_2 \tag{30}$$

Let $T$ be the random variable, which is the earliest time for which the process $\mathbf{X}$ is in a state $x \notin \mathbb{V}$. Let $\widetilde{T}$ be the random variable, which is the earliest time for which the process $\widetilde{\mathbf{X}}$ is in the absorbed state $a$. The formal definitions follow:

$$T = \inf\{t > 0 : (\mathbf{X})_t \notin \mathbb{V}\} \tag{31}$$

$$\widetilde{T} = \inf\{t > 0 : (\widetilde{\mathbf{X}})_t = a\} \tag{32}$$

We refer to $T$ and $\widetilde{T}$ as the exit times.

We start by proving that $T$ and $\widetilde{T}$ are equal in distribution. It will allow us to infer about the time the process $\mathbf{X}$ first exits the set of states $\mathbb{V}$ using process $\widetilde{\mathbf{X}}$. We propose a coupling between processes $\mathbf{X}$ and $\widetilde{\mathbf{X}}$ to show the equality between $T$ and $\widetilde{T}$.

Let $\mathbf{C} = (\mathbf{X}, \widetilde{\mathbf{X}})$ be a coupling of the process and the cut-chain process. Let $\xi_{\mathbf{s}}$ be a sequence of draws from a uniform distribution on $[0,1]$, with elements denoted by $\xi_{s,i}$ and indexed by $i$. Let $\xi_{\mathbf{t}}$ be a sequence of draws from a uniform distribution on $[0,1]$, with elements denoted by $\xi_{t,i}$ and indexed by $i$. Let the initial state of the coupling $C_0$ follow:

$$\begin{aligned}
&X_0 = \widetilde{X}_0 = x \quad\ \text{if } x \in \mathbb{V}, \\
&X_0 = x, \widetilde{X}_0 = a \quad \text{if } x \notin \mathbb{V}.
\end{aligned} \tag{33}$$

Before defining the update rules of $\mathbf{C}$, we define two sets that will be used in the update rules for the coupling. Let $x, y \in \chi$. Let $S_{x,y-}$ and $S_{x,y}$ be the following sets:

$$S_{x,y-} = \{z \in \chi \mid Q(x,y) > 0, z < y\}, \tag{34}$$

$$S_{x,y} = \{z \in \chi \mid Q(x,y) > 0, z \leq y\}. \tag{35}$$

The update rules of $\mathbf{X}$ in the coupling $\mathbf{C}$ follow:

$$X_{i+1} = y \text{ if } \frac{\sum_{z \in S_{x,y-}} Q(x,z)}{\sum_{z \in \chi \setminus \{x\}} Q(x,z)} \leq \xi_{s,i} < \frac{\sum_{z \in S_{x,y}} Q(x,z)}{\sum_{z \in \chi \setminus \{x\}} Q(x,z)}. \tag{36}$$

Note that for both $S_{x,y-}$ and $S_{x,y}$, one has $x \notin S_{x,y-}$ and $x \notin S_{x,y}$ as $Q(x,x) < 0$. Thus, the probability that the process will perform a transition into the same state it was in before is zero.

Let $E_x^{-1}$ be the inverse of the cumulative distribution function (CDF) for the time spent in state $x$. As process $\mathbf{X}$ is a CTMC with generator matrix $Q$, one has that $E^{-1}1_x$ is the CDF of an exponential random variable with parameter $\sum_{z \in \chi \setminus \{x\}} Q(x,z)$. Thus, the time spent in $X_i$ in the coupling $\mathbf{C}$ follows:

$$t_i = t_{i-1} + E_{X_i}^{-1}(\xi_{t,i}). \tag{37}$$

The update rules of $\widetilde{\mathbf{X}}$ in the coupling $\mathbf{C}$ follow:

$$\widetilde{X}_{i+1} = y \text{ if } \frac{\sum_{z \in S_{x,y-}} \widetilde{Q}(x,z)}{\sum_{z \in \mathcal{X} \setminus \{x\}} \widetilde{Q}(x,z)} \leq \xi_{s,i} < \frac{\sum_{z \in S_{x,y}} \widetilde{Q}(x,z)}{\sum_{z \in \mathcal{X} \setminus \{x\}} \widetilde{Q}(x,z)}. \tag{38}$$

Finally, the time spent in $\widetilde{X}_i$ in the coupling $\mathbf{C}$ follows:

$$\widetilde{t}_i = \widetilde{t}_{i-1} + E^{-1}_{\widetilde{X}_i}(\xi_{t,i}). \tag{39}$$

The distinction between time and state coupling is assured by Gillespie [20].

Now we prove a lemma that requires coupling $C$:

**Lemma 1.** *The distributions of $T$ and $\widetilde{T}$ are equal.*

*Proof.* To prove $T \cong \widetilde{T}$, it suffices to prove that under update rules of coupling $\mathbf{C}$, the states of both processes remain equal until process $\mathbf{X}$ reaches a state outside of $\mathbb{V}$ and $\widetilde{\mathbf{X}}$ reaches the absorbing state at the same time.

To formalize it, either Statement (a) or (b) must be true:

(a) There must either exist a $j \in \mathbb{N}$ such that the following statements all hold:

- $X_j \notin \mathbb{V}$
- $\widetilde{X}_j = a$
- For all $i < j$:
$$X_i = \widetilde{X}_i$$
- For all $i < j$:
$$t_i = \widetilde{t}_i \tag{40}$$

(b) Or for all $i$:

$$X_i = \widetilde{X}_i$$
$$t_i = \widetilde{t}_i \tag{41}$$

We will prove this by induction.

We verify for the base case that either the conditions from Statement (a) are satisfied or that $X_0 = \widetilde{X}_0$ at $t = \widetilde{t} = 0$ with $X_0 \in \mathbb{V}$ through a case distinction over the initial states with $x \notin \mathbb{V}$ or not in $x \in \mathbb{V}$:

- **If** $x \notin \mathbb{V}$: From the initial state of the coupling $\mathbf{C}$ in Equation (33), if $X_0 \notin \mathbb{V}$, it implies that $\widetilde{X}_0 = a$. Thus, $X_0 \notin \mathbb{V}$ and $X_0 = a$. It follows that $j = 0$ satisfies the conditions from Equation (40).

- **If** $x \in \mathbb{V}$: From the coupling initial condition in Equation 33, one has $X_0 = \widetilde{X}$ at $t = \widetilde{t} = 0$.

For the induction step, we extend the base case where the conditions from Equation (40) are not satisfied. Let $X_i = \widetilde{X}_i = x$ with $x \in \mathbb{V}$ and $t_{i-1} = \widetilde{t}_{i-1}$. We perform a case distinction based on the location of the next state $X_{i+1} = y$:

- **If** $X_{i+1} = y \in \mathbb{V}$: From the induction step, one has that $\widetilde{X}_i = X_i = x \in \mathbb{V}$. From the assumption that $X_{i+1} = y$, one can conclude that

$$\frac{\sum_{z \in S_{x,y-}} Q(x,z)}{\sum_{z \in \mathcal{X} \setminus \{x\}} Q(x,z)} \leq \xi_{\text{s, i}} < \frac{\sum_{z \in S_{x,y}} Q(x,z)}{\sum_{z \in \mathcal{X} \setminus \{x\}} Q(x,z)} \tag{42}$$

From the definition of $\widetilde{Q}$, if $z \in \mathbb{V}$ and $x \in \mathbb{V}$, one has:

$$\widetilde{Q}(x,z) = Q(x,z) \tag{43}$$

Again, from the definition of $\widetilde{Q}$, one has the following for the absorbing state:

$$\widetilde{Q}(x,a) = \sum_{z\in\chi\backslash\mathbb{V}} Q(x,z) \tag{44}$$

It follows from Equations (44) and (43):

$$\sum_{z\in S_{x,y}} Q(x,z) = \sum_{z\in S_{x,y}\backslash(\chi\backslash\mathbb{V})} Q(x,z) + \sum_{z\in(\chi\backslash\mathbb{V})} Q(x,z) \tag{45}$$

$$\sum_{z\in S_{x,y}} Q(x,z) = \sum_{z\in S_{x,y}\backslash(\chi\backslash\mathbb{V})} \widetilde{Q}(x,z) + \widetilde{Q}(x,a) \tag{46}$$

$$\sum_{z\in S_{x,y}} Q(x,z) = \sum_{z\in S_{x,y}} \widetilde{Q}(x,z) \tag{47}$$

One can repeat the argument in Equation (47) using $S_{x,y-}$ instead of $S_{x,y}$ to conclude equation Equation (47) also holds for $S_{x,y-}$.

From Equations (44) and (43), one can also conclude the following:

$$\sum_{z\in\chi\backslash\{x\}} Q(x,z) = \sum_{z\in\mathbb{V}\backslash\{x\}} Q(x,z) + \sum_{z\in(\chi\backslash\mathbb{V})} Q(x,z) \tag{48}$$

$$\sum_{z\in\chi\backslash\{x\}} Q(x,z) = \sum_{z\in\mathbb{V}\backslash\{x\}} \widetilde{Q}(x,z) + \widetilde{Q}(x,a) \tag{49}$$

$$\sum_{z\in\chi\backslash\{x\}} Q(x,z) = \sum_{z\in\chi\backslash\{x\}} \widetilde{Q}(x,z) \tag{50}$$

It follows from Equations (47) and (50):

$$\frac{\sum_{z\in S_{x,y}} Q(x,z)}{\sum_{z\in\chi\backslash\{x\}} Q(x,z)} = \frac{\sum_{z\in S_{x,y}} \widetilde{Q}(x,z)}{\sum_{z\in\chi\backslash\{x\}} \widetilde{Q}(x,z)} \tag{51}$$

$$\frac{\sum_{z\in S_{x,y-}} Q(x,z)}{\sum_{z\in\chi\backslash\{x\}} Q(x,z)} = \frac{\sum_{z\in S_{x,y-}} \widetilde{Q}(x,z)}{\sum_{z\in\chi\backslash\{x\}} \widetilde{Q}(x,z)} \tag{52}$$

Thus, from the definition of the coupled process in Equation (38), one also has $\widetilde{X}_{i+1} = y$.

For the duration of the transition, from Equation (50), one has $\sum_{z\in\chi\backslash\{x\}} Q(x,z) = \sum_{z\in\chi\backslash\{x\}} \widetilde{Q}(x,z)$, and from the induction hypothesis one has $t_{i-1} = \widetilde{t}_{i-1}$. It follows from Equations (37) and (39) that $t_i = \widetilde{t}_i$ as the parameter for the exponential draw is the same.

- **If** $X_{i+1} = y \notin \mathbb{V}$: Again, from the draw of the coupled process, one has:

$$\frac{\sum_{z\in S_{x,y-}} Q(x,z)}{\sum_{z\in\chi\backslash\{x\}} Q(x,z)} \leq \xi_{\text{s, i}} < \frac{\sum_{z\in S_{x,y}} Q(x,z)}{\sum_{z\in\chi\backslash\{x\}} Q(x,z)} \tag{53}$$

From the chosen order in $\chi$ in Equation 30, as the state $a$ is not contained in $\mathbb{V}$, one can conclude that $a < y$ for $y \in \mathbb{V}$. Further, as the absorbing state is the only state of the process $\mathbf{X}$ which is not in $\mathbb{V}$, it is the lowest possible state reachable by $\widetilde{\mathbf{X}}$. From the choice of the complete order in $\chi$ and the definition of $S_{x,y-}$, one can conclude that the set $S_{x,a^-}$ is empty for every $x \in \chi$. It follows:

$$\sum_{z \in S_{x,a^-}} Q(x,z) = 0 \tag{54}$$

Equation (54) and the update rules from the Coupled process imply:

$$\widetilde{X}_{i+1} = a \text{ if } 0 \leq \xi_{\text{s, i}} < \frac{\widetilde{Q}(x,a)}{\sum_{z \in \chi \setminus \{x\}} \widetilde{Q}(x,z)} \tag{55}$$

From the chosen order in $\chi$ in Equation 30, as $y \notin \mathbb{V}$, for any $w \in \mathbb{V}$, one can conclude that $y < w$. Further, from the definition of $S_{x,y}$, for any state $z \in S_{x,y}$, one has $z \leq y < w$ and $z$ must also not be in $\mathbb{V}$. It follows that:

$$S_{x,y} \subset (\chi \setminus \mathbb{V}) \tag{56}$$

It follows from Equation (56) and the definition of $\widetilde{Q}(x,a)$:

$$\sum_{z \in S_{x,y}} Q(x,z) \leq \sum_{z \in (\chi \setminus \mathbb{V})} Q(x,z) = \widetilde{Q}(x,a) \tag{57}$$

As from Equation (50), the denominator from the quotients from the process and the cut-off process update rule is equal, one can use Equations (50), (53) and (57) for the following:

$$0 \leq \xi_{\text{s, i}} < \frac{\sum_{z \in M_{x,y}} Q(x,z)}{\sum_{z \in \chi \setminus \{x\}} Q(x,z)} \leq \frac{\widetilde{Q}(x,a)}{\sum_{\widetilde{y} \in \chi \setminus \{x\}} \widetilde{Q}(\widetilde{x},\widetilde{y})} \tag{58}$$

Thus, from Equation (55), one has $\widetilde{X}_{i+1} = a$.

For the duration of the transition, again from Equation (50), one has $\sum_{z \in \chi \setminus \{x\}} Q(x,z) = \sum_{z \in \chi \setminus \{x\}} \widetilde{Q}(x,z)$, and from the induction hypothesis $t_{i-1} = \widetilde{t}_{i-1}$, one has $t_i = \widetilde{t}_i$. It follows from Equations (37) and (39) that $t_i = \widetilde{t}_i$ as the parameter for the exponential draw is the same. Furthermore, the index $i$ satisfies the conditions from Statement (a)

It follows from the induction step that either the process can leave to a state outside of $\mathbb{V}$ at the same time and Statement (a) is satisfied, or both processes can remain in $\mathbb{V}$ indefinitely and Statement (b) is satisfied. Thus, one has $T \cong \widetilde{T}$, and the proof is concluded. □

The following Corollary can also arise from the Lemma 1:

**Corollary 1.** *One has* $\mathbb{P}_d(\mathbf{X}(t) = x \mid T > t) = \mathbb{P}_d(\widetilde{\mathbf{X}}(t) = x \mid \widetilde{T} > t)$.

It comes directly from the proof in Lemma 1, where both processes have equal states and times when updated with the coupling rules under the condition that $T > t$ and $\widetilde{T} > t$.

We aim to show that $\widetilde{\mathbf{X}}$ has a quasi-stationary distribution and that it converges to a QSD under the conditional probability that the absorbing state is never reached. To do so, we will propose a third Markov process ${}^a\widetilde{\mathbf{X}}$ created from $\widetilde{\mathbf{X}}$ with the absorbing state removed. We then use the stationary distribution ${}^a\widetilde{\mathbf{X}}$ to find the quasi-stationary distribution of $\widetilde{\mathbf{X}}$ and prove its convergence to it.

The third Markov process ${}^a\widetilde{\mathbf{X}}$, with states $({}^a\widetilde{X}_0, {}^a\widetilde{X}_1, {}^a\widetilde{X}_2, ....)$ in the state space $\mathbb{V} \cup \{a\}$, is defined repeating the notation rules used for defining $\widetilde{\mathbf{X}}$ from $\mathbf{X}$. Let $x, y \in \mathbb{V}$, and $a$ be the absorbing state of $\widetilde{\mathbf{X}}$. The generator matrix ${}^a\widetilde{Q}$ of process ${}^a\widetilde{\mathbf{X}}$ is defined in the following way:

$$\begin{aligned} {}^a\widetilde{Q}(x,y) &= \widetilde{Q}(x,y) \quad \text{if } y \in \mathbb{V} \\ {}^a\widetilde{Q}(x,a) &= 0 \end{aligned} \tag{59}$$

Therefore, process ${}^a\widetilde{\mathbf{X}}$ cannot reach the absorbing state.

We now prove the following:

**Lemma 2.** *For every state $x \in \mathbb{V} \cup \{a\}$, one has:*

$$\mathbb{P}_d({}^a\widetilde{\mathbf{X}}(t) = x) = \mathbb{P}_d(\widetilde{\mathbf{X}}(t) = x \mid T > t) \tag{60}$$

*Furthermore, if $s_d$ is a stationary distribution for ${}^a\widetilde{\mathbf{X}}$, the distribution $s_d$ is also a quasi-stationary distribution for $\widetilde{\mathbf{X}}$.*

*Proof.* As a Markov process under the condition that some states can not be reached is still a Markov process, process $\widetilde{\mathbf{X}}$ under the condition that $T > t$ is a Markov process. Note $Q'$ the generator matrix of a process $\widetilde{\mathbf{X}}$ under the condition that $T > t$.

Let $x, y \in \mathbb{V}$. For any continuous-time Markov process, the transition rate $Q(x, y)$ is defined as ([46]):

$$\widetilde{Q}(x, y) = \lim_{dt \to 0} \frac{\mathbb{P}(\widetilde{X}_{t+dt} = y \mid \widetilde{X}_t = x)}{dt}. \tag{61}$$

Similarly, the transition rate $Q'$ given that the process has not left the state space $\mathbb{V}$ by time $t$ is:

$$Q'(x, y) = \lim_{dt \to 0} \frac{\mathbb{P}(\widetilde{X}_{t+dt} = y \mid \widetilde{X}_t = x \mid T > t)}{dt}. \tag{62}$$

Since the transition from state $x \in \mathbb{V}$ to state $y \in \mathbb{V}$ is independent of the event $T > t$, we have:

$$\begin{aligned} Q'(x, y) &= \lim_{dt \to 0} \frac{\mathbb{P}(\widetilde{X}_{t+dt} = y \mid \widetilde{X}_t = x \mid T > t)}{dt} \\ Q'(x, y) &= \lim_{dt \to 0} \frac{\mathbb{P}(\widetilde{X}_{t+dt} = y \mid \widetilde{X}_t = x)}{dt} \\ Q'(x, y) &= \widetilde{Q}(x, y). \end{aligned} \tag{63}$$

It is left to prove that $Q'(x, a) = 0$ for the absorbing state $a$ to prove the equality between probabilities in the statement of this lemma.

For every state $x \in \mathbb{V}$, the Master Equation ([21]) applied to process $\widetilde{\mathbf{X}}$ under the condition that $T > t$ yields the following:

$$\frac{d}{dt}\mathbb{P}_d(\widetilde{\mathbf{X}}(t) = x \mid T > t) = \sum_{y \in \chi \setminus \{x\}} Q'(y, x)\mathbb{P}_d(\widetilde{\mathbf{X}}(t) = y \mid T > t) - \sum_{z \in \chi \setminus \{x\}} Q'(x, z)\mathbb{P}_d(\widetilde{\mathbf{X}}(t) = x \mid T > t) \tag{64}$$

The Master Equation for the absorbing state $a$ yields:

$$\frac{d}{dt}\mathbb{P}_d(\widetilde{\mathbf{X}}(t) = a \mid T > t) = \sum_{y \in \mathbb{V}} Q'(y, a)\mathbb{P}_d(\widetilde{\mathbf{X}}(t) = y \mid T > t) - \sum_{z \in \mathbb{V}} Q'(a, z)\mathbb{P}_{d_v}(\widetilde{\mathbf{X}}(t) = x \mid T > t) \tag{65}$$

As $a$ is an absorbing state, one has $Q'(a, z) = 0$ for $z \in \mathbb{V}$. It follows:

$$\frac{d}{dt}\mathbb{P}_d(\widetilde{\mathbf{X}}(t) = a \mid T > t) = \sum_{y \in \mathbb{V}} Q'(y, a)\mathbb{P}_d(\widetilde{\mathbf{X}}(t) = y \mid T > t) \tag{66}$$

Under the conditional event that $T > t$, the probability of $\widetilde{\mathbf{X}}$ being absorbed must be zero, and the following must hold for the derivative:

$$\frac{d}{dt}\mathbb{P}_d(\widetilde{\mathbf{X}}(t) = a \mid T > t) = 0 = \sum_{y \in \mathbb{V}} Q'(y, a)\mathbb{P}_d(\widetilde{\mathbf{X}}(t) = y \mid T > t) \tag{67}$$

For all $x \in \mathbb{V}$, for all $z \in \chi$, let $d_x$ be the following initial distribution:

$$\begin{aligned} d_x(z) &= \mathbb{P}(\widetilde{\mathbf{X}}(0) = z \mid T > t) = 0 \text{ If } z \neq x \\ d_x(z) &= \mathbb{P}(\widetilde{\mathbf{X}}(0) = z \mid T > t) = 1 \text{ If } z = x \end{aligned} \tag{68}$$

Equation (67) must be true for every initial distribution $d$ concentrated on $\mathbb{V}$. Thus, by applying Equation (67) for all distributions $d_x$ with $x \in \mathbb{V}$ from Equation (68) yields:

$$\begin{aligned}
0 &= \sum_{y \in \mathbb{V}} Q'(y,a)\mathbb{P}_{d_x}(\widetilde{\mathbf{X}}(t) = y \mid T > t) \\
0 &= Q'(x,a)\mathbb{P}_{d_x}(\widetilde{\mathbf{X}}(t) = x \mid T > t) \\
0 &= Q'(x,a) \qquad (69)
\end{aligned}$$

It means that given the condition that $T > t$, one must have $Q'(x,a) = 0$ for all $x \in \mathbb{V}$.

For all $x, y \in \chi$, it follows from the definition of process ${}^a\widetilde{\mathbf{X}}$, Equation (63), and Equation (69):

$$Q'(x,y) = {}^a\widetilde{Q}(x,y) \qquad (70)$$

From Equation (70), as the process $\widetilde{\mathbf{X}}$ under the condition $T > t$ and the process ${}^a\widetilde{\mathbf{X}}$ have the same generator matrix. For all $x \in \mathbb{V} \cup \{a\}$, one has the following equations:

$$\mathbb{P}_d(\widetilde{\mathbf{X}}(t) = x \mid T > t) = \mathbb{P}_d({}^a\widetilde{\mathbf{X}}(t) = x) \qquad (71)$$

$$\frac{d}{dt}\mathbb{P}_d(\widetilde{\mathbf{X}}(t) = x \mid T > t) = \frac{d}{dt}\mathbb{P}_d({}^a\widetilde{\mathbf{X}}(t) = x) \qquad (72)$$

Let $s_d$ be a stationary probability distribution for process ${}^a\widetilde{\mathbf{X}}$ where $s_d(x)$ is the probability of finding process ${}^a\widetilde{\mathbf{X}}$ at state $x$ when this process is in its stationary distribution $s_d$.

From the definition of a stationary distribution, the distribution $s_d$ is a stationary distribution, if and only if, the following holds for all $t \in \mathbb{R}$ and for $x \in \mathbb{V} \cup \{a\}$:

$$\frac{d}{dt}s_d(x) = \frac{d}{dt}\mathbb{P}_{s_d}({}^a\widetilde{\mathbf{X}}(t) = x) = 0 \qquad (73)$$

From the definition of a quasi-stationary distribution, a distribution $d_\pi$ is a quasi-stationary distribution for $\widetilde{\mathbf{X}}$, if and only if, the following holds for all $t \in \mathbb{R}$ and $y \in \mathbb{V} \cup \{a\}$:

$$\frac{d}{dt}\mathbb{P}_{d_\pi}(\widetilde{\mathbf{X}}(t) = y \mid T > t) = 0 \qquad (74)$$

We now prove that $s_d$ satisfies the requirements for being a QSD distribution in Equation 74 through a case distinction for the states $y \in \mathbb{V} \cup \{a\}$:

- If $y \in \mathbb{V}$: We have the following equality from Equations (72) and (73):

$$\frac{d}{dt}s_d(y) = \frac{d}{dt}\mathbb{P}_{s_d}({}^a\widetilde{\mathbf{X}}(t) = x) = \frac{d}{dt}\mathbb{P}_{s_d}(\widetilde{\mathbf{X}}(t) = y \mid T > t) = 0 \qquad (75)$$

- If $y = a$: For all $t \in \mathbb{R}$, the process ${}^a\widetilde{\mathbf{X}}(t)$ has

$$\frac{d}{dt}\mathbb{P}_{s_d}({}^a\widetilde{\mathbf{X}}(t) = a) = 0 \qquad (76)$$

  as, from its definition, it cannot reach or exit the absorbing state $a$. Therefore, from Equation (72):

$$\frac{d}{dt}\mathbb{P}_{s_d}({}^a\widetilde{\mathbf{X}}(t) = a) = \frac{d}{dt}\mathbb{P}_{s_d}(\widetilde{\mathbf{X}}(t) = a \mid T > t) = 0 \qquad (77)$$

It follows that the distribution $s_d$ satisfies the conditions to be a quasi-stationary distribution for process $\widetilde{X}$ and the proof is concluded. □

As a positive recurrent continuous-time Markov process converges to a stationary distribution, we now prove process ${}^a\widetilde{\mathbf{X}}$ is positive recurrent to prove $\widetilde{\mathbf{X}}$ converges to the same distribution.

The proof below is folklore (See Chapter 2 ([24]) for another proof). We prove it again in this work for the sake of completeness.

**Lemma 3.** *Process ${}^a\widetilde{\mathbf{X}}$ is positive recurrent.*

*Proof.* From the definition of a process $\widetilde{\mathbf{X}}$, the set $\mathbb{V}$ is connected for $\widetilde{\mathbf{X}}$. It follows from the definition of ${}^a\widetilde{\mathbf{X}}$ that set $\mathbb{V}$ is also connected for ${}^a\widetilde{\mathbf{X}}$.

It follows from the connectivity the states in $\mathbb{V}$ for process ${}^a\widetilde{\mathbf{X}}$ and that $\mathbb{V}$ is finite, that for every two states $x, y \in \mathbb{V}$ there exists finite sequence states $A_{x,y} = ({}^a\widetilde{X}_0 = x, {}^a\widetilde{X}_1, ...., {}^a\widetilde{X}_{n_a-1} = y)$ of length $n_A$ of non-zero probability of being executed by process ${}^a\widetilde{\mathbf{X}}$. Furthermore, the probability of execution does not asymptotically approach zero as the sequence $A_{x,y}$ is finite.

Let $\delta t \in \mathbb{R}^+_{>0}$. Let $E_x$ be an exponential random variable of parameter $\sum_{y\in\mathcal{X}} {}^aQ(x, y)$. The probability noted by:

$$\mathbb{P}_{\delta t}(x, y) = \mathbb{P}({}^a\widetilde{X}_0 = x, {}^a\widetilde{X}_1, ...., {}^a\widetilde{X}_{n_A-1} = y, \sum_{i=0}^{n_A-1} E_{{}^a\widetilde{X}_i} < \delta t) \tag{78}$$

is the probability of process ${}^a\widetilde{\mathbf{X}}$ executing the sequence $A_{x,y}$ in less than $\delta t$ time. As the sequence $({}^a\widetilde{X}_1 = x, {}^a\widetilde{X}_1, ...., {}^a\widetilde{X}_{n_s} = y)$ has a probability non-zero of being executed by the process, it does not asymptotically approach zero, and the sum $\sum_{i=0}^{n_A-1} E_{{}^a\widetilde{X}_i}$ is finite, one can conclude that there exists a constant $p_A \in \mathbb{R}^+$ for every $x, y \in \mathbb{V}$ such that:

$$\mathbb{P}_{\delta t}(x, y) \geq p_A > 0 \tag{79}$$

Note the probability that the process has been in state $y$ starting from $x$ in under $\delta t$ time as the following:

$${}^\delta J(x, y) = \mathbb{P}(\exists t \in (0, \delta t] \text{ such that } \mathbf{X}(t) = y \mid \mathbf{X}(0) = x) \tag{80}$$

For every two states $x, y \in \mathbb{V}$, the probability that the process has been in state $y$ starting from $x$ in under $\delta t$ time is at least the probability of reaching state $y$ from $x$ using the sequence states $S_{x,y}$. Therefore, with Equation (79), we can conclude:

$${}^\delta J(x, y) \geq \mathbb{P}_{\delta t}(x, y) \geq p_A > 0 \tag{81}$$

Thus, for every two states $x, y \in \mathbb{V}$, the probability ${}^\delta J(x, y)$ is at least $p_A$.

Let $\tau_x$ be the random variable regarding the time process ${}^a\widetilde{\mathbf{X}}$ firstly returns to $x$:

$$\tau_x = \min\{t \in \mathbb{R}^+ \mid ({}^a\widetilde{\mathbf{X}})_t = x, ({}^a\widetilde{\mathbf{X}})_0 = x\} \tag{82}$$

Further, let $\delta\tau_x$ be the following random variable:

$$\delta\tau_x = \min\{i\delta t \in \mathbb{R}^+ \mid ({}^a\widetilde{\mathbf{X}})_t = x, i \in \mathbb{N}^+, i\delta t \geq t, ({}^a\widetilde{\mathbf{X}})_0 = x\} \tag{83}$$

which is the first time multiple of $\delta t$ for which the process has already been in state $x$.

For any execution of process ${}^a\widetilde{\mathbf{X}}$, we perform a case distinction:

- **If $\tau_x = t_e$ and $t_e$ is a multiple of $\delta t$:** Then one has $\tau_x = \delta\tau_x$
- **If $\tau_x = t_e$ and $t_e$ is not a multiple of $\delta t$:** Then one has $\tau_x < \delta\tau_x$

Consequently:

$$\mathbb{E}(\tau_x) \leq \mathbb{E}(\delta\tau_x) \tag{84}$$

The probability of $\tau_x$ being greater than a given time $i\delta t$ with $i \in \mathbb{N}$ is given by the probability that process has never been in state $x$ apart from its initial state:

$$\mathbb{P}(\delta\tau_x > (i+1)\delta t) = \sum_{y\in\mathbb{V}, y\neq x} \mathbb{P}(X_i = y, X_j \neq x \mid 0 \leq j \leq i, \leq X_0 = x) \tag{85}$$

For $(i+1)\delta t$, one can consider Markov memoryless property to write that the probability of not reaching state $x$ until time $(i+1)\delta t$ is 1 minus that probability of reaching $x$ from $y$ from in under $\delta t$ time times $\mathbb{P}(\delta\tau_x > i\delta t)$:

$$\mathbb{P}(\delta\tau_x > (i+1)\delta t) = \sum_{y\in\mathbb{V}, y\neq x} \mathbb{P}(X_i = y, X_j \neq x \mid 0 \leq j \leq i, X_0 = x) \cdot (1 - {}^{\delta}J(x,y)) \tag{86}$$

Further, Equation (81) implies $p_A \leq {}^{\delta}J(x,y)$ for all $x, y \in \mathbb{V}$ which yields:

$$\begin{aligned} &\mathbb{P}(\tau_x > (i+1)\delta t) \\ &\leq \sum_{y\in\mathbb{V}, y\neq x} \mathbb{P}(X_i = y, X_j \neq x \mid 0 \leq j \leq i, X_0 = x) \cdot (1 - p_A) \\ &\leq (1 - p_A) \cdot \sum_{y\in\mathbb{V}, y\neq x} \mathbb{P}(X_i = y, X_j \neq x \mid 0 \leq j \leq i, X_0 = x) \end{aligned} \tag{87}$$

Using Equation (85) in Equation (87) yields:

$$\mathbb{P}(\delta\tau_x > (i+1)\delta t) \leq (1 - p_A)\mathbb{P}(\delta\tau_x > i\delta t) \tag{88}$$

It yields by recurrence for $n \in \mathbb{N}$:

$$\mathbb{P}(\delta\tau_x > (i+k)\delta t) \leq (1 - p_A)^k \mathbb{P}(\delta\tau_x > i\delta t) \tag{89}$$

From Equation (81), one has that $p_A > 0$. Thus, let $i_d \in \mathbb{N}, i_d > 0$, we can bound the expected value of $\tau_x$ in the following manner:

$$\mathbb{E}(\delta\tau_x) = \sum_{i=0}^{\infty} \mathbb{P}(\delta\tau_x > i\delta t) \tag{90}$$

$$\mathbb{E}(\delta\tau_x) = \sum_{i=0}^{i_d} \mathbb{P}(\delta\tau_x > i\delta t) + \sum_{i=i_d+1}^{\infty} \mathbb{P}(\delta\tau_x > i\delta t) \tag{91}$$

Using Equation (89) yields the following:

$$\mathbb{E}(\delta\tau_x) \leq \sum_{i=0}^{i_d} \mathbb{P}(\delta\tau_x > i\delta t) + \mathbb{P}(\delta\tau_x > i_d\delta t) \sum_{i=i_d+1}^{\infty} (1 - p_A)^{i - i_d} \tag{92}$$

As $\sum_{i=0}^{i_d} \mathbb{P}(\delta\tau_x > i\delta t)$ and $\mathbb{P}(\delta\tau_x > i_d\delta t)$ are finite, and $\sum_{i=i_d+1}^{\infty} (1-p_A)^{i-i_d}$ converges to finite value the following holds:

$$\mathbb{E}(\delta\tau_x) < \infty \tag{93}$$

From Equation (84), it finally yields:

$$\mathbb{E}(\tau_x) \leq \mathbb{E}(\delta\tau_x) < \infty \tag{94}$$

Since this expected value for the return time is finite, state $x$ is positive recurrent, which implies process ${}^{a}\widetilde{\mathbf{X}}$ is positive recurrent. □

Lemma (3) proves process ${}^{a}\widetilde{\mathbf{X}}$ is positive recurrent. It is known that a connected positive recurrent continuous-time Markov process converges to a unique stationary distribution ([8]), which allows us to prove the following lemma.

**Lemma 4.** *Let $d_{\mathbb{V}}$ be a initial distribution concentrated on $\mathbb{V}$. A quasi-stationary distribution of process $\widetilde{\mathbf{X}}$ is given by the following unique limit:*

$$\lim_{t\to\infty} \mathbb{P}_{d_{\mathbb{V}}}(\widetilde{\mathbf{X}}(t) = x \mid \widetilde{T} > t) = \lim_{t\to\infty} \mathbb{P}_{d_{\mathbb{V}}}(\mathbf{X}(t) = x \mid T > t) = \pi_q(x) \tag{95}$$

*Proof.* As Lemma 3 assures process ${}^{a}\widetilde{\mathbf{X}}$ is positive recurrent, process ${}^{a}\widetilde{\mathbf{X}}$ is a continuous-time Markov process, it follows from ([8]) that process ${}^{a}\widetilde{\mathbf{X}}$ converges to a stationary distribution $s_d$ given by:

$$\lim_{t\to\infty} \mathbb{P}_{d_{\mathbb{V}}}({}^{a}\widetilde{\mathbf{X}}(t) = x) = s_d(x) \tag{96}$$

Furthermore, as the set of states $\mathbb{V}$ is connected in process ${}^{a}\widetilde{\mathbf{X}}$, process ${}^{a}\widetilde{\mathbf{X}}$ cannot reach the absorbing state $a$, and the initial distribution $d_{\mathbb{V}}$ is concentrated on $\mathbb{V}$, the distribution given by Equation (96) is unique for all $d_{\mathbb{V}}$.

The equality of distributions from Lemma 2, yields the following:

$$\lim_{t\to\infty} \mathbb{P}_{d_{\mathbb{V}}}({}^{a}\widetilde{\mathbf{X}}(t) = x) = \lim_{t\to\infty} \mathbb{P}_{d_{\mathbb{V}}}(\widetilde{\mathbf{X}}(t) = x \mid \widetilde{T} > t) = s_d(x) \tag{97}$$

Finally, the equality from Corollary 1 yields:

$$\lim_{t\to\infty} \mathbb{P}_{d_{\mathbb{V}}}(\widetilde{\mathbf{X}}(t) = x \mid \widetilde{T} > t) = \lim_{t\to\infty} \mathbb{P}_{d_{\mathbb{V}}}(\mathbf{X}(t) = x \mid T > t) = s_d(x) \tag{98}$$

From Lemma 2, as $s_d$ is a stationary distribution for ${}^{a}\widetilde{\mathbf{X}}$, one has $s_d$ is a quasi-stationary distribution for $\widetilde{\mathbf{X}}$. Thus, let $\pi_q = s_d$ to conclude the proof. □

Van et al. ([43]), proved the following theorem:

**Theorem 3.** *Let $\mathcal{S} = \bigcup_{k=1}^{L} \mathcal{S}_k$ be the union of the set of communicating classes of a continuous-time Markov process $\mathbb{X}$ of generator matrix $Q$. Let $\alpha_k$ be the eigenvalue of the communicating class $S_k$. Let $-x$ be an eigenvalue. Let $P_E$ be the matrix exponential of $Q$.*

*Let $\mathbf{u}$ represent a probability distribution over $\mathcal{S}$. Then the following statements are equivalent:*

*(a)* $\mathbf{u}$ *is a quasi-stationary distribution for* $\mathbb{X}$

*(b)* $\mathbf{u}$ *is a quasi-stationary distribution for* $\mathbb{X}$

*(c)* $\mathbf{u}$ *is $x$-invariant for $Q$ for some $x > 0$.*

*(d)* $\mathbf{u}$ *is $x$-invariant for $P_E$ for some $x > 0$.*

*(e)* $\mathbf{u}$ *is $\alpha_k$-invariant for $Q$ for some $k \in 1, 2, ...L$.*

*(f)* $\mathbf{u}$ *is $\alpha_k$-invariant for $P$ for some $k \in 1, 2, ...L$.*

Their proof strategy involves showing the equivalence of various properties of a Markov process using its generator matrix $Q$ and the matrix exponential $P_E$. It begins by summing over the transient states and demonstrating that $x = 0$ contradicts the transient nature of the states. It then shows that an $x$-invariant distribution for $Q$ is also $x$-invariant for $P_E$, and vice versa, by taking derivatives and letting $t \to 0$. The proof proceeds to establish the equivalence of different statements (b) through (e) and demonstrates that a quasi-stationary distribution $u$ leads to $u$ being $x$-invariant for $Q$. It concludes by breaking down $u$ and $Q$ into block-triangular forms and applying known results to show that $x$ must equal one of the eigenvalues $a_k$.

We now rewrite Corollary 2 from ([43]) derived from Theorem 3 to adapt it to this work:

**Corollary 2.** *The quasi-stationary distribution $\pi_q$ for process $\widetilde{\mathbf{X}}(t)$ yields:*

$$\mathbb{P}_{\pi_q}(\widetilde{T} > t) = \exp(-\lambda t) \tag{99}$$

*where $\lambda$ is the negative of the non-zero eigenvalue with the maximal real part of $\widetilde{Q}$.*

The parameter $\lambda$ is called the decay parameter.

From Lemma 2 and Lemma 1, we get the following collolary:

**Corollary 3.** *One has that $\mathbb{P}_{\pi_q}(T > t) = \mathbb{P}_{\pi_q}(\widetilde{T} > t) = \exp(-\lambda t)$*

#### B.2 End of Theorem 1 Proof

To construct the proof of Theorem 1, we wish to use the quasi-stationary distribution of process $\widetilde{\mathbf{X}}$ as a bound for $\mathbb{P}_d(T > t)$. For this, we will use the jump chain $J$ matrix derived from the generator matrix $\widetilde{Q}$. One has that $J(x, y)$ gives the probability of the state $\widetilde{X}_i$ being $y$ if the previous state was $\widetilde{X}_{i-1} = x$. Formally, let $J(x, y)$ with $x \in \mathbb{V}$ and $y \in \mathbb{V} \cup \{a\}$ be the probability given by:

$$\begin{aligned} J(x,y) &= \frac{\widetilde{Q}(x,y)}{\sum_{z\in\chi\setminus\{x\}} \widetilde{Q}(x,z)} \quad \text{if } x \neq y \text{ and } x \in \mathbb{V} \\ J(x,x) &= 1 \quad \text{if} \sum_{y\in\chi\setminus\{x\}} Q(x,y) = 0 \\ J(x,x) &= 0 \quad \text{otherwise} \end{aligned} \tag{100}$$

A lemma necessary for constructing the bound using the jump chain follows:

**Lemma 5.** *For all $x \in \chi$, one has:*

$$\mathbb{P}(\widetilde{T} \leq t \mid \widetilde{X}_0 = x) \leq \sum_{z\in\chi} J(x,z)\mathbb{P}(\widetilde{T} \leq t \mid \widetilde{X}_0 = z) \tag{101}$$

*Proof.* One can prove this by case distinction.

**If** $x \in \mathbb{V}$:

Let the set $\Omega_x$ be the set of finite sequences $\mathbf{S^x}$ of states in $\chi$ starting at state $x \in \mathbb{V}$ and ending at the absorbing state $a$. Let the elements of these sequences of index $i$ be given by $S_i^x$ and the length of each of these sequences be given by $n_x$.

The probability that the hitting time $\widetilde{T}$ is smaller than $t$ can be written as the sum of the probability of all sequences in $\mathbf{S}^x$ starting in $x$ being absorbed before time $t$:

$$\mathbb{P}(\widetilde{T} \leq t \mid \widetilde{X}_0 = x) = \sum_{\mathbf{S^x}\in\Omega_x} \mathbb{P}(\forall i, 0 \leq i < n_x, X_i = S_i^x \wedge \sum_{i=0}^{n_x-1} E_{S_i^x} \leq t \mid \widetilde{X}_0 = x) \tag{102}$$

Thus, through the substitution using the jump chain, one has:

$$\mathbb{P}(\widetilde{T} \leq t \mid \widetilde{X}_0 = x) = \sum_{\mathbf{S^x}\in\Omega_x} \prod_{i=0}^{n_x-1} J(S_i^x, S_{i+1}^x) \cdot \mathbb{P}(\sum_{i=0}^{n_x-1} E_{S_i^x} \leq t) \tag{103}$$

Let $\Omega_{x,y}$ be the the set of sequences $\mathbf{S^{x,y}}$ starting at state $x \in \mathbb{V}$ and whose first transition after leaving state $x$ is to $y \in \chi$. Let the elements of these sequences of index $i$ be given by $S_i^{x,y}$, and the length of each of those sequences be given by $n_{x,y}$.

As $x \in \mathbb{V}$, one has that $x \neq a$ as $a \notin \mathbb{V}$. If state $x$ is not the absorbing one and every sequence $\Omega_x$ is absorbed, there must be at least one transition to $x$ to any other state. This means there must be a $z \in \chi$ with $J(x, z) > 0$. One can also conclude that any sequence in $\Omega_x$ must be an element of $\bigcup_{z\in\chi} \Omega_{x,z}$, so $\Omega_x \subset \bigcup_{z\in\chi} \Omega_{x,z}$. Further, every sequence $\mathbf{S^{x,z}}$ in $\bigcup_{z\in\chi} \Omega_{x,z}$ starts in $x$ and is absorbed, which implies $\mathbf{S^{x,z}} \in \Omega_x$ and $\bigcup_{z\in\chi} \Omega_{x,z} \subset \Omega_x$. It follows $\bigcup_{z\in\chi} \Omega_{x,z} = \Omega_x$.

Thus, one has:

$$\mathbb{P}(\widetilde{T} \leq t \mid \widetilde{X}_0 = x) = \sum_{\mathbf{S^x}\in\bigcup_{z\in\chi}\Omega_{x,z}} \prod_{i=0}^{n_x-1} J(S_i^x, S_{i+1}^x) \cdot \mathbb{P}(\sum_{i=0}^{n_x-1} E_{S_i^x} \leq t) \tag{104}$$

From the definition of $\Omega_{x,z}$, there can only be one transition after leaving state $x$ for the first time. Thus, for all $z, z' \in \chi$ one has that $\Omega_{x,z} \cap \Omega_{x,z'} = \emptyset$, and we can break the sum in Equation 104 in the following manner for every state $y \in \chi$:

$$\mathbb{P}(\widetilde{T} \leq t \mid \widetilde{X}_0 = x) = \sum_{\mathbf{S^x} \in \bigcup_{z \in \chi \setminus \{y\}} \Omega_{x,z}} \prod_{i=0}^{n_x - 1} J(S_i^x, S_{i+1}^x) \cdot \mathbb{P}(\sum_{i=0}^{n_x-1} E_{S_i^x} \leq t)$$
$$+ \sum_{\mathbf{S^{x,y}} \in \Omega_{x,y}} \prod_{i=0}^{n_{x,y}-1} J(S_i^{x,y}, S_{i+1}^{x,y}) \cdot \mathbb{P}(\sum_{i=0}^{n_{x,y}-1} E_{S_i^{x,y}} \leq t) \tag{105}$$

One can use this to analyze the terms of the sum independently. Note the sum $P_y$ as the following:

$$P_y = \sum_{\mathbf{S^{x,y}} \in \Omega_{x,y}} \prod_{i=0}^{n_{x,y}-1} J(S_i^{x,y}, S_{i+1}^{x,y}) \cdot \mathbb{P}(\sum_{i=0}^{n_{x,y}-1} E_{S_i^{x,y}} \leq t) \tag{106}$$

From the definition of $\Omega_{x,y}$, one has $S_0^{x,y} = x$ and $S_1^{x,y} = y$, as the sequence of states starts at $x$ and its first transition is to $y$. Thus, for all sequences in $\Omega_{x,y}$ one has as a common factor the first transition probability $J(x, y)$:

$$P_y = J(x,y) \left( \sum_{\mathbf{S^{x,y}} \in \Omega_{x,y}} \prod_{i>0}^{n_{x,y}-1} J(S_i^{x,y}, S_{i+1}^{x,y}) \cdot \mathbb{P}(\sum_{i=0}^{n_{x,y}-1} E_{S_i^{x,y}} \leq t) \right) \tag{107}$$

Further, as $E_{S_i^{x,y}}$ is an exponentially distributed positive random variable, for the term $\mathbb{P}(\sum_{i=0}^{n_{x,y}-1} E_{S_i^{x,y}} \leq t)$, one can bound the probability by removing the exponential random variable $E_{S_0^{x,y}}$ regarding the time spent on the first state:

$$\mathbb{P}\left(\sum_{i=0}^{n_{x,y}-1} E_{S_i^{x,y}} \leq t\right) \leq \mathbb{P}\left(\sum_{i>0}^{n_{x,y}-1} E_{S_i^{x,y}} \leq t\right) \Rightarrow \tag{108}$$

$$\sum_{\mathbf{S^{x,y}} \in \Omega_{x,y}} \prod_{i>0}^{n_{x,y}-1} J(S_i^{x,y}, S_{i+1}^{x,y}) \cdot \mathbb{P}\left(\sum_{i=0}^{n_{x,y}-1} E_{S_i^{x,y}} \leq t\right) \leq \sum_{\mathbf{S^{x,y}} \in \Omega_{x,y}} \prod_{i>0}^{n_{x,y}-1} J(S_i^{x,y}, S_{i+1}^{x,y}) \cdot \mathbb{P}\left(\sum_{i>0}^{n_{x,y}-1} E_{S_i^{x,y}} \leq t\right) \tag{109}$$

Let $\mathbf{S}_d$ be a subsequence from the set $\Omega_{x,y}$ that had the first element removed. The first state of $\mathbf{S}_d$ is $y$, and it is absorbed since it is in $\Omega_{x,y}$. Thus, from the definition of $\Omega_y$, the sequence $\mathbf{S}_d$ is in $\Omega_y$. Furthermore, take any sequence from $\mathbf{S}_p$ from $\Omega_y$. A new sequence $\mathbf{S}_r$ with $x$ as the first element and the elements from $\mathbf{S}_p$ following is in $\Omega_{x,y}$. Consequently:

$$\sum_{\mathbf{S^{x,y}} \in \Omega_{x,y}} \prod_{i>0}^{n_{x,y}-1} J(S_i^{x,y}, S_{i+1}^{x,y}) \cdot \mathbb{P}\left(\sum_{i>0}^{n_{x,y}-1} E_{S_i^{x,y}} \leq t\right) = \sum_{\mathbf{S^{x,y}} \in \Omega_y} \prod_{i=0}^{n_y-1} J(S_i^{x,y}, S_{i+1}^{x,y}) \cdot \mathbb{P}\left(\sum_{i=0}^{n_y-1} E_{S_i^{x,y}} \leq t\right) \tag{110}$$

It then follows from Equation (102):

$$\mathbb{P}(\widetilde{T} \leq t \mid \widetilde{X}_0 = y) = \sum_{\mathbf{S^{x,y}} \in \Omega_y} \prod_{i=0}^{n_y-1} J(S_i^{x,y}, S_{i+1}^{x,y}) \cdot \mathbb{P}\left(\sum_{i=0}^{n_y-1} E_{S_i^{x,y}} \leq t)\right) \tag{111}$$

And finally for ${}^sP_y$ from Equations (107) and (109):

$$P_y \leq J(x,y)\mathbb{P}(T \leq t \mid \widetilde{X}_0 = y) \tag{112}$$

One can then use Equations (105) and (112) to all $z \in \chi$ to conclude the following:

$$\mathbb{P}(\widetilde{T} \leq t \mid \widetilde{X}_0 = x) = \sum_{z \in \chi} P_z$$
$$\mathbb{P}(\widetilde{T} \leq t \mid \widetilde{X}_0 = x) \leq \sum_{z \in \chi} J(x,z)\mathbb{P}(\widetilde{T} \leq t \mid \widetilde{X}_0 = z) \tag{113}$$

This concludes the part of the proof for $x \in \mathbb{V}$.

**If** $x \notin \mathbb{V}$:

If $x \notin \mathbb{V}$, one can conclude that:

$$P(\widetilde{T} \leq t \mid X_0 = x) = 1 \tag{114}$$

Further, as $J$ is a jump chain matrix, the sum of its rows is equal to 1, which yields:

$$\sum_{z \in \chi} J(x, z) = 1 \tag{115}$$

Therefore:

$$P(\widetilde{T} \leq t \mid X_0 = x) = 1 \tag{116}$$

$$\sum_{z \in \chi} J(x, z) P(\widetilde{T} \leq t \mid X_0 = x) = 1 \tag{117}$$

and the proof is concluded.

□

From the equality of conditional probabilities of Corollary 1 and Lemma 102, we get the following:

**Corollary 4.** *For $x \in \chi$, one has:*

$$\mathbb{P}(T \leq t \mid X_0 = x) \leq \sum_{z \in \chi} J(x, z) \mathbb{P}(T \leq t \mid X_0 = z) \tag{118}$$

From Corolary 3, for theorem 1, we wish to find $A$ such that the following holds:

$$\mathbb{P}_d(T > t) \geq \mathbb{P}_{\pi_q}(T > t) - A \tag{119}$$

$$\mathbb{P}_d(T > t) \geq \exp(-\lambda t) - A \tag{120}$$

We will use Corollary 4 to prove that the steps in Algorithm 1 yield an $A$ for which Equation (119) holds.

Let $\epsilon \in \mathbb{R}$. Let calculate_jump_chain be a function that returns the jump chain according to Equation (100). Let max_loops $\in \mathbb{N}$. Bellow one can find Algorithm 1 that bounds $\mathbb{P}_d(T > t)$ for any distribution using the decay parameter and the constant $A$:

Let $q_i, D_i, D'_i, A_i$ be the arrays resulting from Algorithm's 1 respective variables $q, D, D', A$ after the execution of the for loop $i$ times.

Let us define the functions of time $\hat{q}_i$, $\hat{D}_i$ and $\hat{D}'_i$ from $\mathbb{R}^+$ to $\mathbb{R}$ using the arrays from Algorithm's 1. Their definitions follow:

$$\hat{q}_i(t) = \sum_{x \in \chi} q_i[x] \mathbb{P}(T \leq t \mid X_0 = x) \tag{121}$$

$$\hat{D}_i(t) = \sum_{x \in \chi} D_i[x] \mathbb{P}(T \leq t \mid X_0 = x) \tag{122}$$

$$\hat{D}'_i(t) = \sum_{x \in \chi} D'_i[x] \mathbb{P}(T \leq t \mid X_0 = x) \tag{123}$$

Now, we prove a lemma regarding an inequality presented in all steps of the algorithm:

**Lemma 6.** *For any $i$, for any time $t$, one has that:*

$$\hat{D}_i(t) - \hat{q}_i(t) \geq \mathbb{P}_d(T \leq t) - \mathbb{P}_{\pi_q}(T \leq t) \tag{124}$$

*Proof.* One can prove the lemma by induction. The base case comes directly from the initialization of $\hat{D}_0$ and $\hat{q}_0$, which are initialized with the distribution $d$ and the quasi-stationary distribution $q$. Thus:

$$\hat{q}_0(t) = \sum_{x\in\chi} q_0[x]\mathbb{P}(T \le t \mid X_0 = x) = \mathbb{P}_{\pi_q}(T \le t)$$
$$\hat{D}_0(t) = \sum_{x\in\chi} D_0[x]\mathbb{P}(T \le t \mid X_0 = x) = \mathbb{P}_d(T \le t)$$

and

$$\hat{D}_0(t) - \hat{q}_0(t) = \mathbb{P}_d(T \le t) - \mathbb{P}_{\pi_q}(T \le t) \tag{125}$$

Let $x \in \mathbb{V} \cup \{a\}$. For the induction step, one can perform a case distinction between the states for which $q_i[x] \ge D_i[x]$ and the states for which $q_i[x] < D_i[x]$:

- **If for a state $x \in \chi$ one has $q_i[x] \ge D_i[x]$:** Then according to Algorithm 1 $q_{i+1}[x]$ and $D_{i+1}[x]$ are:

$$\begin{aligned}&\mathbb{P}(T \le t \mid X_0 = x)D_i[x] - \mathbb{P}(T \le t \mid X_0 = x)q_i[x]\\ &= (D_i[x] - q_i[x]) \cdot \mathbb{P}(T \le t \mid X_0 = x)\\ &= -q_{i+1}[x] \cdot \mathbb{P}(T \le t \mid X_0 = x)\end{aligned} \tag{126}$$

  and

$$D_i'[x] = 0 \tag{127}$$

- **If for a state $x \in \chi$ one has $q_i[x] < D_i[x]$:** Then according to Algorithm 1 $D_i'[x]$ and $q_{i+1}[x]$ are:

$$\begin{aligned}&\mathbb{P}(T \le t \mid X_0 = x)D_i[x] - \mathbb{P}(T \le t \mid X_0 = x)q_i[x]\\ &= (D_i[x] - q_i[x]) \cdot \mathbb{P}(T \le t \mid X_0 = x)\\ &= D_i'[x] \cdot \mathbb{P}(T \le t \mid X_0 = x)\end{aligned} \tag{128}$$

  and

$$q_{i+1}[x] = 0 \tag{129}$$

From Corollary 4, one has:

$$D_i'[x] \cdot \mathbb{P}(T \le t \mid X_0 = x) \le \sum_{z\in\chi} J(x,z)D_i'[x]\mathbb{P}(T \le t \mid X_0 = z) \tag{130}$$

Equation (130) implies:

$$\sum_{x\in\chi}\sum_{z\in\chi} J(x,z)D_i'[x]\mathbb{P}(T \le t \mid X_0 = z) \ge \sum_{x\in\chi} D_{i+1}'[x]\mathbb{P}(T \le t \mid X_0 = z) \tag{131}$$

From Algorithm (1) line 21 and the definition of $\hat{D}_{i+1}$, one has:

$$\sum_{x\in\chi} D_{i+1}[x]\mathbb{P}(T \le t \mid X_0 = z) = \sum_{x\in\chi}\sum_{z\in\chi} J(x,z)D_i'[x]\mathbb{P}(T \le t \mid X_0 = z) \tag{132}$$

It is:

$$\begin{aligned}&\hat{D}_i(t) - \hat{q}_i(t)\\ &= \sum_{x\in\chi} D_i[x]\mathbb{P}(T \le t \mid X_0 = x) - \sum_{x\in\chi} q_i[x]\mathbb{P}(T \le t \mid X_0 = x)\\ &= \sum_{x\in\chi} \mathbb{P}(T \le t \mid X_0 = x)(D_i[x] - q_i[x])\end{aligned} \tag{133}$$

From the case distinction, for every state $x$ the result from the subtraction is either $D'_i[x]$ or $q_{i+1}[x]$ which results in:

$$\hat{D}_i(t) - \hat{q}_i(t) = \sum_{x \in \mathcal{X}} D'_i[x]\mathbb{P}(T \leq t \mid X_0 = x) - \sum_{x \in \mathcal{X}} q_{i+1}[x]\mathbb{P}(T \leq t \mid X_0 = x) \tag{134}$$

From Equations (130), (131), and (132), one has:

$$\begin{aligned}
&\hat{D}_i(t) - \hat{q}_i(t) \\
&\leq \sum_{x \in \mathcal{X}} \sum_{z \in \mathcal{X}} D'_i[x]J(x,z)\mathbb{P}(T \leq t \mid X_0 = z) - q_{i+1}[x]\mathbb{P}(T \leq t \mid X_0 = x) \\
&= \sum_{x \in \mathcal{X}} D_{i+1}[x]\mathbb{P}(T \leq t \mid X_0 = x) - q_{i+1}[x]\mathbb{P}(T \leq t \mid X_0 = x) \\
&= \hat{D}_{i+1}(t) - \hat{q}_{i+1}(t)
\end{aligned} \tag{135}$$

Thus, one has:

$$\hat{D}_{i+1}(t) - \hat{q}_{i+1}(t) \geq \hat{D}_i(t) - \hat{q}_i(t) \tag{136}$$

This concludes the proof. □

Bellow, we prove the final lemma to validate Algorithm 1.

**Lemma 7.** *Let $A_i$ be $A$ from Algorithm 1 after $i$ rounds of execution. One can conclude the following:*

- $\mathbb{P}_d(T > t) \geq \exp(-\lambda t) - A_i$
- $A_i \geq A_{i+1}$

*Proof.* From the definition of $A_i$, as $q_i[x]$ is positive, and as for all $x$ probabilities follow $\mathbb{P}(T \leq t \mid X_0 = x) \leq 1$, one has:

$$\begin{aligned}
&A_i \\
&= \sum_{x \in \mathcal{X}} D_i[x] \\
&\geq \sum_{x \in \mathcal{X}} D_i[x] - q_i[x] \\
&\geq \sum_{x \in \mathcal{X}} D_i[x]\mathbb{P}(T \leq t \mid X_0 = x) - q_i[x]P(T \leq t \mid X_0 = x) \\
&\geq \hat{D}_i(t) - \hat{q}_i(t)
\end{aligned} \tag{137}$$

Thus from Lemma 6, one also has the following:

$$\begin{aligned}
&A_i \\
&\geq \mathbb{P}_d(T \leq t) - \mathbb{P}_{\pi_q}(T \leq t) \\
&\geq (1 - \mathbb{P}_d(T > t)) - (1 - \mathbb{P}_{\pi_q}(T > t)) \\
&\geq -\mathbb{P}_d(T > t)) + \mathbb{P}_{\pi_q}(T > t)
\end{aligned} \tag{138}$$

Thus,

$$\mathbb{P}_d(T > t) \geq \mathbb{P}_{\pi_q}(T > t) - A_i \tag{139}$$

From Corollary 2 and Equation (139), one has:

$$\mathbb{P}_d(T > t) \geq \mathbb{P}_{\pi_q}(T > t) - A_i = \exp(-\lambda t) - A_i \tag{140}$$

Therefore, the first statement from this lemma holds.

It is left to prove that $A_i \geq A_{i+1}$. To prove this, one can perform a case distinction between the states for which $q_i[x] \geq D_i[x]$ and the states for which $q_i[x] < D_i[x]$.

- **If for a state $x \in \chi$ one has $q_i[x] \geq D_i[x]$:** Then according to Algorithm 1 for this state:

$$D_i[x] - q_i[x] = -q_{i+1}[x] \tag{141}$$

  Furthermore, from Algorithm 1, for this state $D'_{i+1}[x] = 0$. Thus, one has $D'_i[x] \leq D_i[x]$.

- **If for a state $x \in \chi$ one has $q_i[x] \leq D_i[x]$:** Then according to Algorithm 1 for this state:

$$D_i(x) - q_i[x] = D'_i[x] \tag{142}$$

  Thus, as $q_i[x]$ is positive, one can conclude that $D'_i[x] \leq D_i[x]$

From the case distinction, one can conclude that for every $x$, one has:

$$D'_i[x] \leq D_i[x] \tag{143}$$

It follows:

$$\sum_{x\in\chi} D'_i[x] \leq \sum_{x\in\chi} D_i[x] \tag{144}$$

Finally, as $J$ is a probability jump chain, the sum of the elements in each of the rows of $J$ is one. It also implies the following:

$$\sum_{x\in\chi} JD'_i[x] = \sum_{x\in\chi} D'_i[x] \tag{145}$$

Algorithm 1 line 21 implies the following:

$$\sum_{x\in\chi} D_{i+1}[x] = \sum_{x\in\chi} JD'_i[x] \tag{146}$$

From Equations (144),(146) and (145), one has:

$$A_i = \sum_{x\in\chi} D_i[x] \geq \sum_{x\in\chi} D'_i[x] = \sum_{x\in\chi} JD'_i[x] = \sum_{x\in\chi} D_{i+1}[x] = A_{i+1} \tag{147}$$

The proof is concluded. □

Lemma 7 concluded the prove of Theorem 1.

##### B.3 Proof of Proposition 1

In this section, we apply further restrictions to the process $\mathbf{X}$ and $\widetilde{\mathbf{X}}$. We assume that $\mathbf{X}$ is a birth and death process on state space $\chi = \mathbb{N}$, and consequently, the cut-off process $\widetilde{\mathbf{X}}$ is also a birth and death chain. Without any loss of generality, we assume the absorbed state from the cut-off process $\widetilde{\mathbf{X}}$ follows $a < x$ for any state $x \in \chi$.

We wish to find the set of distributions $\beta$ such that if $d \in \beta$ one has that $\mathbb{P}_d(T > t)$ is at least $\mathbb{P}_{\pi_q}(T > t)$ for all $t \in \mathbb{R}$, allowing us to bound the exit probability of the distributions on the set $\beta$ using the decay parameter of $\widetilde{\mathbf{X}}$. Differently than Theorem 1, here, a constant $A$ is not required for the bound. We refer to distributions from $\beta$ as quasi-stationary bounded distributions.

As a point of departure, we prove a stronger bound relation than the one proposed in Lemma 5, which uses the assumption that $\widetilde{\mathbf{X}}$ is a birth-death process.

**Lemma 8.** *For any two states $y, x \in \mathbb{V}$, with $y < x$, one has:*

$$\mathbb{P}(\widetilde{T} \leq t \mid \widetilde{X}_0 = x) \leq \mathbb{P}(\widetilde{T} \leq t \mid \widetilde{X}_0 = y) \tag{148}$$

*Proof.* Let the set $\Omega_x$ be the set of finite sequences $\mathbf{S^x}$ starting at state $x \in \mathbb{V}$ and ending at the absorbing state $a$. Let the elements of these sequences of index $i$ be given by $S_i^x$, and the length of each of those sequences be given by $n_S$.

For any state $x \in \mathbb{V}$, the probability that the hitting time $T$ is smaller than $t$ can be written as all the state sequences that end at state $a$ and the process is able to execute this state sequence before time $t$:

$$\mathbb{P}(\widetilde{T} \leq t \mid \widetilde{X}_0 = x) = \sum_{\mathbf{S^x} \in \Omega_x} \mathbb{P}(\forall i, 0 \leq i < n_S, X_i = S_i^x \wedge \sum_{i=0}^{n_S-1} E_{S_i^x} \leq t \mid \widetilde{X}_0 = x) \tag{149}$$

Thus, through the substitution using the jump chain, one has:

$$\mathbb{P}(\widetilde{T} \leq t \mid \widetilde{X}_0 = x) = \sum_{\mathbf{S^x} \in \Omega_x} \prod_{i=0}^{n_S-1} J(S_i^x, S_{i+1}^x) \cdot \mathbb{P}\left(\sum_{i=0}^{n_S-1} E_{S_i^x} \leq t\right) \tag{150}$$

For any sequence $\mathbf{S^x} \in \Omega_x$, note the finite sub-sequence $\mathbf{R^x}$ as the sub-sequence whose elements are equal to the elements in $\mathbf{S^x}$ until the state $y \in \mathbb{V}$ firstly appears in the sequence, with $y < x$, and the length of each sequence is given by $n_R$. One can note the set of these corresponding sub-sequences as $\Gamma_x$. The definition follows:

$$\mathbf{R^x} = \{R_i^x \mid R_i^x = S_i^x \wedge i < \min\{j : S_j = y\}\} \tag{151}$$

Further, for any sequence $\mathbf{S^x} \in \Omega_x$, note the sub-sequence $\mathbf{L^x}$ as the sequence containing all elements from $\mathbf{S^x}$ after the element $y$ firstly appears in the sequence inclusively with $y < x$ and the length of each sequence is $n_L$. One can note the set of these corresponding sub-sequences as $\Lambda_x$. The definition follows:

$$\mathbf{L^x} = \{L_i^x \mid L_i^x = S_i^x \wedge i \geq \min\{j : S_j = y\}\} \tag{152}$$

Both $\mathbf{R^x}$ and $\mathbf{L^x}$ are well defined, as for any $y < x$, as the process $\widetilde{\mathbf{X}}$ is a birth-and-death process and all sequences in $\Omega_x$ are absorbed with $a \leq y < x$, there must be at least one element of the sequence $S_j^x$ for which $S_j^x = y$.

For a sequence $\mathbf{S^x} \in \Omega_x$, from the definition of $\mathbf{R^x}$ and $\mathbf{L^x}$, there is a non-empty sequence $\mathbf{R^x} \in \Gamma_x$ and a corresponding sequence $\mathbf{L^x}$ with the remaining elements of $\mathbf{S^x}$ in $\Lambda_x$. One can then rewrite $\mathbb{P}(T \leq t \mid \widetilde{X}_0 = x)$ in the following manner using sequences $\mathbf{L^x}$ and $\mathbf{R^x}$:

$$\mathbb{P}(\widetilde{T} \leq t \mid \widetilde{X}_0 = x) = \sum_{\mathbf{L^x} \in \Lambda_x} \prod_{i=0}^{n_L-1} p_{L_i^x, L_{i+1}^x} \left(\sum_{\mathbf{R^x} \in \Gamma x} \prod_{j=0}^{n_R-1} p_{R_j^x, R_{j+1}^x} \cdot \mathbb{P}\left(\sum_{S_i^x \in \mathbf{S^x}} E_{S_i^x} \leq t\right)\right) \tag{153}$$

As $E_{S_i^x}$ is a positive random variable, one can bound the following term as follows:

$$\mathbb{P}\left(\sum_{i=0}^{n_S-1} E_{S_i^x} \leq t\right) = \mathbb{P}\left(\sum_{i=0}^{n_L-1} E_{L_i^x} + \sum_{i=0}^{n_R-1} E_{R_i^x} \leq t\right) \tag{154}$$

$$\mathbb{P}\left(\sum_{i=0}^{n_S-1} E_{S_i^x} \leq t\right) \leq \mathbb{P}\left(\sum_{i=0}^{n_L-1} E_{L_i^x} \leq t\right) \tag{155}$$

Thus, it follows:

$$\mathbb{P}(\widetilde{T} \leq t \mid \widetilde{X}_0 = x) \leq \sum_{\mathbf{L^x} \in \Lambda_x} \prod_{i=0}^{n_L-1} p_{L_i^x, L_{i+1}^x} \left(\sum_{\mathbf{R^x} \in \Gamma x} \prod_{i=0}^{n_R-1} p_{R_i^x, R_{i+1}^x} \cdot \mathbb{P}\left(\sum_{i=0}^{n_L-1} E_{L_i^x} \leq t\right)\right) \tag{156}$$

$$\mathbb{P}(\widetilde{T} \leq t \mid \widetilde{X}_0 = x) \leq \sum_{\mathbf{L^x} \in \Lambda_x} \prod_{i=0}^{n_L-1} p_{L_i^x, L_{i+1}^x} \cdot \mathbb{P}\left(\sum_{i=0}^{n_L-1} E_{L_i^x} \leq t\right) \left(\sum_{\mathbf{R^x} \in \Gamma x} \prod_{i=0}^{n_R-1} p_{R_i^x, R_{i+1}^x}\right) \tag{157}$$

As $\sum_{\mathbf{R^x}\in\Gamma x}\left(\prod_{R_i^x\in\mathbf{R^x}} p_{R_i^x,R_{i+1}^x}\right)$, is the probability reaching state $y$ from state $x$ for the first time, it can be bounded by 1. Thus, one has:

$$\mathbb{P}(\widetilde{T}\leq t \mid \widetilde{X}_0 = x) \leq \sum_{\mathbf{L^x}\in\Lambda_x}\prod_{i=0}^{n_L-1} p_{L_i^x,L_{i+1}^x}\cdot\mathbb{P}\left(\sum_{i=0}^{n_L-1} E_{L_i^x}\leq t\right) \tag{158}$$

Equation (150) applied to state $y$ yields the following:

$$\mathbb{P}(\widetilde{T}\leq t \mid \widetilde{X}_0 = x) \leq \sum_{\mathbf{L^x}\in\Lambda_x}\prod_{i=0}^{n_L-1} p_{L_i^x,L_{i+1}^x}\cdot\mathbb{P}\left(\sum_{i=0}^{n_L-1} E_{L_i^x}\leq t\right) = \mathbb{P}(\widetilde{T}\leq t \mid \widetilde{X}_0 = y) \tag{159}$$

The proof is then concluded. □

From Corollary 1, we extend Lemma 8 for process $\mathbb{X}$:

**Corollary 5.** *For any two states $y, x \in \mathbb{V}$, with $y < x$, one has:*

$$\mathbb{P}(T\leq t \mid X_0 = x) \leq \mathbb{P}(T\leq t \mid X_0 = y) \tag{160}$$

With Lemma 8, we define a sufficient condition for an initial distribution to be quasi-stationary bounded.

**Theorem 4.** *Let $\widetilde{X}$ be a continuous time birth-death Markov process with an absorbing state $a$ in state space $\mathbb{V}\cup\{a\}\subset\mathbb{N}$. Without loss of generality, let $a < x$ for every state $x\in\mathbb{V}$. Let $\widetilde{J}$ be the jump chain from process $\widetilde{X}$. Let $\widetilde{T}$ be the random variable representing the time until absorption of process $\widetilde{X}$. Let $d_{\mathbb{V}}$ be a distribution concentrated on $\mathbb{V}$.*

*Let $\pi_q$ be the quasi-stationary distribution given by:*

$$\lim_{t\to\infty}\mathbb{P}_{d_{\mathbb{V}}}(\widetilde{\mathbf{X}}(t) = x \mid \widetilde{T} > t) = \pi_q(x) \tag{161}$$

*For all $t\in\mathbb{R}$, the initial distribution $d$ of $\mathbf{X}$ follows:*

$$\mathbb{P}_d(\widetilde{T} > t) \geq \mathbb{P}_{\pi_q}(\widetilde{T} > t) \tag{162}$$

*If the statement below holds:*

- *For every state $y\in\chi$, there exists a function $v_y : \chi\to\mathbb{R}^+$:*

$$\sum_{w\leq y, y\in\chi} v_y(w) \geq d(y) \tag{163}$$

*Such that the sum of all functions $v_y$ for every state $x$ follows:*

$$\pi_q(x) \geq \sum_{y\in\chi} v_y(x) \tag{164}$$

*Where $\pi_q(x)$ is the probability measure of the process being found state $x$ in the quasi-stationary distribution.*

*Proof.* Equation (95) assures the QSD $\pi_q$ is well-defined.

From the law of total probability, one has for distributions $d$ and $\pi_q$:

$$\mathbb{P}_d(\widetilde{T}\leq t) = \sum_{x\in\mathbb{V}}\mathbb{P}(\widetilde{T}\leq t \mid \widetilde{X}_0 = x)d(x) \tag{165}$$

For each state $y\in\chi$ and from the Lemma assumption for $v_y$, one has:

$$\begin{gathered}\sum_{w\leq y, y\in\chi} v_y(w) \geq d(y) \Rightarrow \\ \mathbb{P}(\widetilde{T}\leq t \mid \widetilde{X}_0 = y)\sum_{w\leq y, y\in\chi} v_y(w) \geq \mathbb{P}_d(\widetilde{T}\leq t \mid \widetilde{X}_0 = y)d(y) \Rightarrow \\ \sum_{w\leq y, y\in\chi}\mathbb{P}(\widetilde{T}\leq t \mid \widetilde{X}_0 = y)v_y(w) \geq \mathbb{P}_d(\widetilde{T}\leq t \mid \widetilde{X}_0 = y)d(y)\end{gathered} \tag{166}$$

Lemma 8 assures us that $\mathbb{P}(\widetilde{T} \leq t \mid \widetilde{X}_0 = w) \geq \mathbb{P}(T \leq t \mid \widetilde{X}_0 = y)$ as $w \leq y$. It follows:

$$\sum_{w \leq y, y \in \chi} \mathbb{P}(T \leq t \mid \widetilde{X}_0 = w) v_y(w) \geq \mathbb{P}_d(T \leq t \mid \widetilde{X}_0 = y) d(y) \tag{167}$$

As Equation (167) applies for all $y \in \chi$, by summing over all states, one has:

$$\begin{aligned}
&\mathbb{P}_d(T \leq t) \\
&= \sum_{x \in \mathbb{V}} \mathbb{P}_d(T \leq t \mid \widetilde{X}_0 = x) d(x) \\
&\leq \sum_{y \in \mathbb{V}} \sum_{w \leq y, y \in \chi} \mathbb{P}(T \leq t \mid \widetilde{X}_0 = w) v_y(w) \\
&= \sum_{w \in \mathbb{V}} \mathbb{P}(T \leq t \mid \widetilde{X}_0 = w) \cdot (\sum_{y \in \mathbb{V}} v_y(w)) \\
&\leq \sum_{w \in \mathbb{V}} \mathbb{P}(T \leq t \mid \widetilde{X}_0 = w) \pi_q(w) \\
&= \mathbb{P}_{\pi_q}(T \leq t)
\end{aligned} \tag{168}$$

As $\mathbb{P}(T \leq t) = 1 - \mathbb{P}(T > t)$, from Equation (168), one has:

$$\mathbb{P}_d(T > t) \geq \mathbb{P}_{\pi_q}(T > t) \tag{169}$$

This concludes the proof □

Lemma 1 allows one to extend Theorem 4 to the original process. Furthermore, it allows us to deduce an important property for bounded birth and death chains.

**Lemma 9.** *Let* $\mathbf{X}$ *be a birth-and-death continuous-time Markov whose state space is* $\{0, 1, ...., P\} = \chi \subseteq \mathbb{N}$ *whose generator matrix is* $Q$.

*Let* $a \in \mathbb{N}$ *and* $a < P$. *Let* $\mathbb{V} = \{a+1, a+2, ...., P\}$. *Let* $y \in \mathbb{V} \cup \{a\}$ *Let* $\widetilde{\mathbf{X}}$ *be a birth death Markov process whose state space is* $\mathbb{V} \cup \{a\}$ *, the generator matrix of* $\widetilde{\mathbf{X}}$ *is:*

$$\begin{aligned}
&\widetilde{Q}(x, y) = Q(x, y) \quad \text{if } x \in \mathbb{V} \\
&\widetilde{Q}(a, y) = 0
\end{aligned} \tag{170}$$

*Let* $T$ *be the random variable corresponding to the time process* $\mathbf{X}$ *reaches a state outside of* $\mathbb{V}$. *Let* $\widetilde{T}$ *be the random variable corresponding to the time process* $\widetilde{\mathbf{X}}$ *reaches its absorbing state* $a$.

*Let* $d_P$ *be a distribution concentrated on state* $P$. *It follows for all* $t \in \mathbb{R}$:

$$\mathbb{P}_{d_P}(T > t) \geq \mathbb{P}_{\pi_q}(\widetilde{T} > t) \tag{171}$$

*Proof.* For this proof, one can use Theorem 4.

As required by Theorem 4, one must propose a function for each state $x \in \chi$ and verify it follows the inequality in the theorem statement.

- For state $P$: $v_P = \pi_q$, where $\pi_q$ is the quasi-stationary distribution of $\widetilde{\mathbf{X}}$ given by Lemma 4 whose existence is assured by the fact that the state space of $\widetilde{\mathbf{X}}$ is connected.
- For $x \in \chi$, $x \neq B$: $v_x = 0$

We now verify that the proposed functions satisfy the two conditions from Theorem 4.

Firstly, we perform a case distinction for states $B$ and $x \in \chi, x \neq B$ for the first condition:

- **State** $P$: For state $P$, with $v_P = \pi_q$:

$$d(P) = 1 \tag{172}$$

As $\pi_q$ is a distribution:

$$\sum_{y \in \chi} \pi_q(y) = 1 \tag{173}$$

As $y \leq P$ for all $y \in \chi$, from Theorem 4 requirements, one has:

$$\sum_{y \leq P} \pi_q(y) = \sum_{y \in \chi} \pi_q(y) = 1 \tag{174}$$

It follows that:

$$\sum_{y \leq P} \pi_q(y) = d(P) = 1 \tag{175}$$

So the function $v_P$ satisfies Theorem 4's condition for state $P$.

- **State** $x \in \chi, x \neq P$: For this state, one has:

$$d(x) = 0 = v_x(x) \tag{176}$$

which also satisfies Theorem 4's condition for state $x$.

It follows that the functions $v_x$ and $v_P$ satisfy the first condition.

Secondly, Since $v_B = \pi_q$ and $v_x = 0$ for $x \in \chi, x \neq B$, one can conclude that:

$$\sum_{y \in \chi} v_y = \pi_q \tag{177}$$

It follows that the functions $v_x$ and $v_P$ satisfy the second condition.

The proof is concluded. □

Lemma 9 concludes the proof of Proposition 1 without loss of generality.
